# The landscape of fitness effects of putatively functional noncoding mutations in humans

**DOI:** 10.1101/2025.05.14.654124

**Authors:** Chenlu Di, Swetha Ramesh, Jason Ernst, Kirk E. Lohmueller

## Abstract

While annotations of noncoding regions in the human genome are increasing, the fitness effects of mutations in these regions remain unclear. Here, we leverage these functional genomic annotations and human polymorphism data to infer the distributions of fitness effects of new noncoding mutations in humans. Our novel approach controls for mutation rate variation and linked selection along the genome. We find distinct patterns of selection in putative enhancers, promoters, and conserved noncoding regions. While mutations in enhancers are often neutral, approximately 30% of mutations in promoters are deleterious. The most conserved noncoding regions, showing reduced divergence across mammals and primates, have the highest proportion of deleterious mutations. Notably, while we infer the most conserved sites across mammals and primates are enriched for deleterious mutations, such conserved sites only account for a minority of the deleterious mutations in noncoding regions. For example, the top 5% of conserved noncoding sites encompass fewer than 20% of deleterious mutations, indicating that functional noncoding regions vary widely in the distribution of their evolutionary constraint. Our findings highlight the dynamic evolution of gene regulation and shifting selection pressures over deep evolutionary timescales. Consistent with this finding, we infer mutations in ∼7-9% of the noncoding genome are deleterious. These insights have broad implications for using comparative genomics to identify non-neutrally evolving sequences in the human genome.

## INTRODUCTION

Since King and Wilson (King & Wilson 1975) hypothesized that the genetic variants critical for evolutionary innovation and complex traits would influence gene expression, there has been tremendous interest in studying genetic variation in the noncoding regions of genomes. Indeed, empirical examples of the importance of noncoding regulatory variation in evolution and complex traits continue to grow (Wray 2007; Bickel et al. 2011; Ernst et al. 2011; Wittkopp & Kalay 2011; Gruber et al. 2012; Maurano et al. 2012; Vernot et al. 2012; Connelly et al. 2013; Gusev et al. 2014; Roadmap Epigenomics Consortium et al. 2015; Li et al. 2016; Boyle et al. 2017; Signor & Nuzhdin 2018; Wang et al. 2018). Further, most GWAS hits in humans fall in noncoding regions (Hindorff et al. 2009; Ernst et al. 2011; Maurano et al. 2012; Gusev et al. 2014; Roadmap Epigenomics Consortium et al. 2015). In an effort to better understand the noncoding genome, consortia like ENCODE and Roadmap Epigenomics have annotated features in the human genome that correlate with gene expression and other biological functions. Such studies have now identified millions of putative cis-regulatory elements throughout the genome (The ENCODE Project Consortium 2012; Roadmap Epigenomics Consortium et al. 2015; Meuleman et al. 2020; Moore et al. 2020).

Nevertheless, our understanding of the fitness effects of noncoding mutations lags behind understanding of amino-acid changing mutations. Aided by the ease of functional annotation, the fitness effects of amino acid changes have been extensively studied (Eyre-Walker et al. 2006; Boyko et al. 2008; Bataillon & Bailey 2014; Kim et al. 2017; Zhen et al. 2021) using population genetic methods. Such studies revealed that about 25% of nonsynonymous mutations are strongly deleterious (|*s*|>0.01), 20% are neutral (|*s*|<10^-5^), <1% are beneficial, and the rest are weakly to moderately deleterious (10^-5^<|*s*|<10^−2^). This distribution of fitness effects (DFE), or, the probability distribution of selection coefficients (*s*) of each individual mutation, is a fundamental parameter in evolutionary genetics. Studies of the DFE for noncoding mutations lag further behind, however. One early study (Torgerson et al. 2009) which inferred a DFE for noncoding mutations used a limited dataset, focusing on a subset of human-mouse conserved regions, and only included 35 individuals. While this was a rigorous study when it was published in 2009, it is not clear whether this DFE applies to all noncoding regulatory regions. More recent studies have leveraged functional genomic annotations and deeper polymorphism data to infer fitness effects of noncoding mutations (Arbiza et al. 2013; Gronau et al. 2013; Gulko et al. 2015; Gussow et al. 2017; Vitsios et al. 2021; Dukler et al. 2022), but have been limited by models that only infer a set of selection coefficients, rather than a full DFE. Thus, it is not possible to compare these estimates with those of the DFE for amino acid changing mutations.

As an alternative approach to assessing fitness effects of mutations, comparative genomic methods have revealed mutations in 5-6% of the human genome are likely deleterious (Mouse Genome Sequencing Consortium et al. 2002; Cooper et al. 2005; Siepel et al. 2005; Pollard et al. 2010; Lindblad-Toh et al. 2011). However, these estimates are not without controversy. There is mounting evidence of turnover of functional sites across mammalian evolution, where mutations at a particular position in the genome may have different fitness consequences throughout evolutionary history (Meader et al. 2010; Ponting & Hardison 2011; Ponting et al. 2011; Rands et al. 2014; Huber et al. 2020). Further, another recent comparative genomic study of 240 placental mammal species estimated that while mutations in as much 10.7% of the human genome are deleterious, <40% of those positions could be reliably identified from patterns of constraint (Christmas et al. 2023). Further, we recently developed a score to aggregate functional genomic information to predict whether a base is likely to be under evolutionary constraint (Grujic et al. 2020). We observed depletions in polymorphism in predicted functional sites that showed little evidence of phylogenetic constraint. As such, comparative genomic methods across distantly related species may not reveal selection occurring only on more recent timescales. Thus, there is a need for estimates of fitness effects of noncoding mutations in humans.

Here, we combine functional genomic annotations of the noncoding genome with patterns of genetic variation and population genetic models to estimate the DFE and assess the evolutionary implications of noncoding mutations. Through this analysis, we seek to advance our understanding of the evolutionary forces shaping mutations in the human genome and provide insights into the role of noncoding variation in human health and disease.

## RESULTS

### Patterns of genetic variation vary across noncoding annotations

To examine patterns of genetic variation across different noncoding functional annotations, we used the universal pan-tissue genomic annotations from Vu & Ernst 2022. Genomic regions are classified into different chromatin states defined jointly from the enrichment of epigenomic signals of multiple cell types using ChromHMM (Ernst & Kellis 2012; Vu & Ernst 2022). Intersecting these annotations with polymorphism data from the 1000 Genomes Project, we found that patterns of polymorphism differ substantially across chromatin states (Fig.1a, Table.S1b). Notably, heterochromatin (HET, yellow) and quiescent chromatin (Quies, gray) show higher genetic diversity, while regions marked by promoters (TSS, PromF, BivProm, red) and transcription-related states (Tx, TxEx, TxEnh, green) exhibit lower genetic diversity. Chromatin states with low genetic diversity also have a more negative value of Tajima’s *D*, which indicates excess rare variants compared to the neutral model. Taken together, these results suggest negative selection is acting on mutations in chromatin states in the bottom-left corner (Fig.1a), including promoters and transcription-related states. In contrast, enhancer-associated regions (EnhA, EnhWk) show intermediate genetic diversity and Tajiama’s *D* values. Promoter regions (TSS, PromF, and BivProm) are generally positioned further from inactive chromatin states (Quies and HET) compared to enhancer regions, emphasizing their distinct polymorphism pattern.

**Fig. 1:**
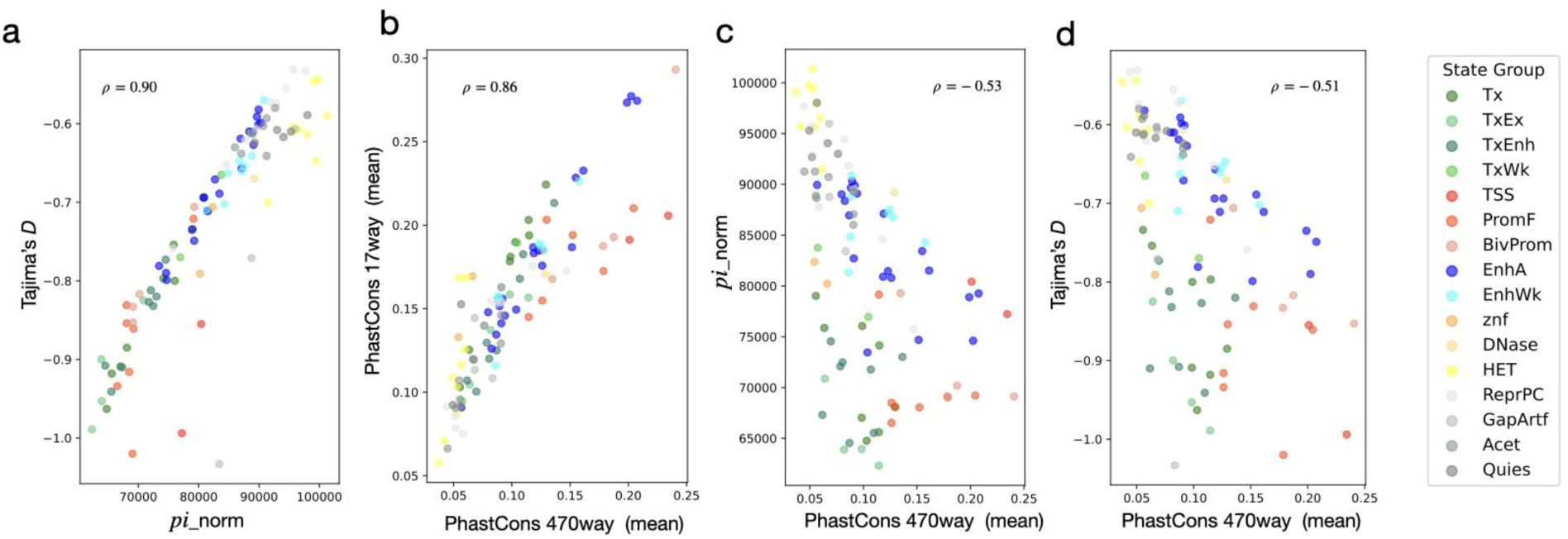
Patterns of genetic variation across chromatin state groups. **a**, Tajima’s *D* is positively correlated with the normalized pairwise nucleotide diversity (*pi*_norm, *pi* normalized by mutation rate and sequence length, see details in Methods:Processing functional genomic annotations) across various chromatin states (Pearson correlation coefficient, *ρ*=0.90). Each point represents a genomic region color-coded by chromatin state group, as indicated in the legend on the right. Chromatin states include transcription (Tx), transcription starting sites (TSS), promoter flanking (PromF), bivalent promoters (BivProm), active enhancers (EnhA), heterochromatin (HET), quiescent (Quies) and others (Table S1c). **b**, PhastCons 17way primate constraint is positively correlated with PhastCons 470way mammalian constraint across chromatin state groups (Pearson correlation coefficient, *ρ*= 0.86). **c**, **d**, *pi*_norm and Tajima’s *D* are weakly correlated with PhastCons 470way scores with a Pearson correlation coefficient, *ρ* = -0.53 and -0.51, respectively.

The degree of phylogenetic constraint also varies across chromatin states (Fig.1b). We quantified constraint in primates using the PhastCons 17way scores and in mammals using the PhastCons 470way conservation scores downloaded from UCSC (Siepel et al. 2005; Perez et al. 2025). Genomic regions that are highly conserved in the 17way alignment also tend to be conserved in the broader 470way alignment (*ρ* = 0.86). Consistent with polymorphism patterns, heterochromatin (HET) and quiescent chromatin (Quies) show lower levels of conservation, clustering toward the lower-left of the plot. In contrast, promoter regions (TSS, PromF, and BivProm) tend to cluster at the upper-right area of the plot, reflecting higher levels of evolutionary constraint. Enhancer regions (EnhA, EnhWk) display greater variation, indicating variable levels of evolutionary constraint. We also found that genetic diversity in the human population negatively correlates with the constraint across species (Fig.1c, Fig.S1c), suggesting some broad similarities in selection across these evolutionary times (though we investigate this relationship in more detail in the section: Mutations in phylogenetically conserved enhancers and promoters are more deleterious). In addition, chromatin states that are more constrained across species are more likely to be enriched with rare variants (Fig.1d, Fig.S1d), again suggesting that constrained genomic regions may be under negative selection in the human population. Further zooming in to enhancer states, we found that three enhancer chromatin states (EnhA1, 3, 9) are clustered at the bottom-left corner with low genetic diversity and are enriched with rare variants (Fig.1a and Fig.S1a). These three chromatin states are enhancers found in most cell types. This result indicates that negative selection may be stronger in general enhancers, especially strong enhancers. These findings suggest that genetic diversity and evolutionary constraint vary significantly across chromatin states, potentially reflecting different levels of natural selection acting on these functional annotations.

Mutation rate variation has been found to confound the inference of natural selection (Castellano et al. 2020; Soni et al. 2024). We found that the average mutation rates vary within and between different chromatin states but are not correlated with Tajima’s *D* (Fig.S1e). This indicates that other evolutionary forces, such as natural selection and non-equilibrium demography, may have a larger influence on the patterns of rare variants compared to mutation rates.

### DFE for enhancers and promoters

To quantify natural selection acting on mutations in enhancers and promoters more precisely, we used a model-based method, Fit*dadi* (Gutenkunst et al. 2009; Kim et al. 2017; Huang et al. 2023), to estimate the distribution of fitness effects (DFE) from patterns of polymorphism as summarized by the site frequency spectrum (SFS). We controlled for demographic history, linked selection, and mutation rate variation along the genome in these inferences (see Methods: DFE inference). We define “promoters” as genomic regions that are “TSS”- the transcription starting sites, “PromF”- the promoters in various tissues, or “BivProm”- the bivalent promoters annotated by ChromHMM, and define “active enhancers” as genomic regions annotated by twenty “EnhA” ChromHMM states. Our method accounts for mutation rate variation along the human genome by using a high-resolution mutation rate map (Seplyarskiy et al. 2022; see Methods: Calculating mutation rates). We also account for variation in linked selection along the genome by inferring demographic models from polymorphism in nearby quiescent sites, which are genomic regions annotated as being in the“Quie” chromatin state by ChromHMM.

Both active enhancers and promoters exhibit a greater proportion of neutral mutations and fewer strongly deleterious mutations compared to nonsynonymous mutations (Fig. 2a). Although the proportion of neutral mutations in enhancers is as high as 91%, we can still confidently reject the neutral model, which assumes that all new mutations are neutral. Specifically, we found that the negative selection model using a gamma DFE fits the SFS significantly better than the neutral model, with a log-likelihood difference exceeding 2255 (p-value < 2.22×10^-308^) (Methods:DFE inference), providing statistical support for the conclusion that mutations in enhancers are not neutrally evolving. Compared to enhancers, promoters have a smaller proportion of neutral mutations and a higher proportion of weakly deleterious mutations (Fig. 2a). Thus, we confidently reject the neutral model for promoters with the gamma DFE showing an improvement in log-likelihood exceeding 4141 (p-value < 2.22×10^-308^) compared to the neutral model. However, compared to non-synonymous mutations, both promoters and enhancers exhibit only a small proportion of moderately and strongly deleterious mutations.

**Fig. 2:**
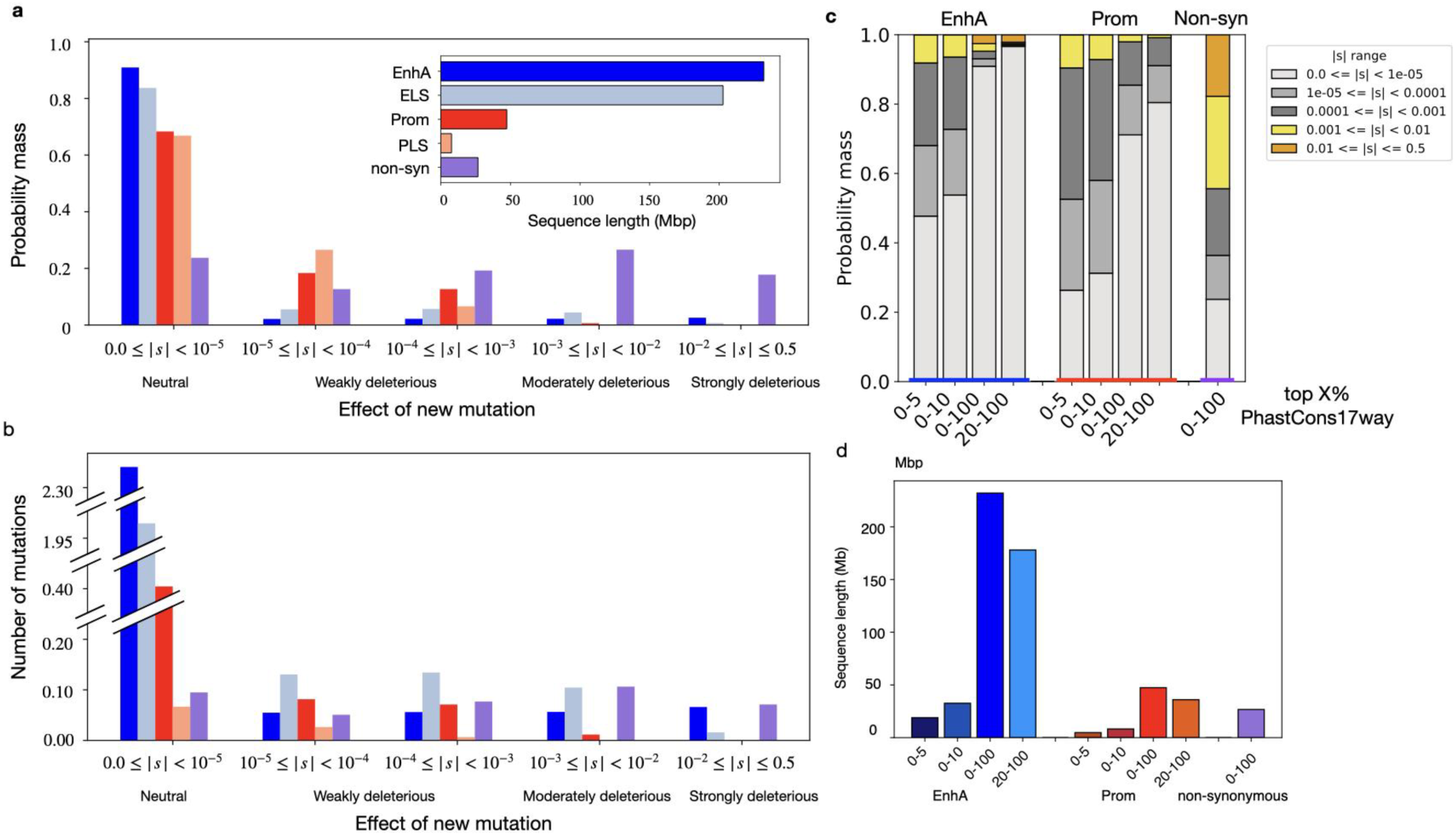
DFEs of mutations in enhancers and promoters. **a**, The probability mass of new mutations across different ranges of selection coefficients (|*s*|) for enhancers, promoters, and non-synonymous mutations. The effect of new mutation is categorized into four classes based on the strength of selection: neutral (0.0≤ |*s*|<10^-5^), weakly deleterious (10^-5^≤ |*s*|<10^-3^), moderately deleterious (10^-3^≤ |*s*| <10^-2^), and strongly deleterious (10^-2^≤ |*s*| <0.5). Active enhancers (EnhA) from ChromHMM annotation are colored by dark blue (the first bar from the left), while light blue bars (the second bar from the left) correspond to enhancers (ELS) annotated by the ENCODE project. Promoters (Prom) based on ChromHMM annotation are shown in red (the third bar from the left), while light red bars (the fourth bar from the left) indicating promoters (PLS) annotated by the ENCODE project. Nonsynonymous mutations (data from Kim et al., 2017) are depicted in purple (the last bar from the left). The insert shows the number of base pairs in each annotation. **b**, The distributions of the expected number of mutations per haploid genome per generation having different effects on fitness in enhancers, promoters, and nonsynonymous sites. The number of mutations (y-axis) is the product of the probability mass of having mutations with a given selection coefficient, the sequence length and the average mutation rate of the group. Groups in (**b**) are indicated using the same colors as in (**a**). **c**, The DFEs of conserved enhancers and promoters. The left and middle groups of bars are active enhancers and promoters annotated by ChromHMM, and the right-most group corresponds to non-synonymous mutations. The x-axis indicates the topX% PhastCons17way scores, the y-axis indicates the probability mass of mutations with different selection coefficients marked by the color. For example, the first left bar shows the DFE of conserved enhancers of which the PhastCons17way score is within top 5%. **d**, The sequence length of conserved active enhancers (EnhA) and promoters (Prom) annotated by ChromHMM. The order is the same as in (c), blue represents active enhancers, red represents promoters and purple represents non-synonymous mutations.

Different samples, cell types, and annotation strategies might influence the identification of enhancers and promoters. To examine the robustness of our findings, we compared the DFEs in active enhancers (EnhA) and promoters (PromF, BivProm, TSS) annotated by ChromHMM (Vu & Ernst, 2022) to candidate cis-regulatory elements with enhancer-like signatures (cCREs-ELS) and promoter-like signatures (cCREs-PLS) identified by ENCODE CREs (ENCODE Cis-Regulatory Elements, The ENCODE Project Consortium 2012; Moore et al. 2020). Reassuringly, the DFEs inferred for the different functional annotations are largely concordant across the annotations and their overlapping regions (Fig.S2). Specifically, enhancers from both annotations consistently exhibit a larger proportion of neutral mutations, whereas promoters show a greater proportion of weakly and moderately deleterious mutations.

We further investigated the DFE of proximal-enhancers (cCRE-pELS) that are enhancers (cCREs-ELS) located within 2kb of transcription start sites and distal-enhancers (cCRE-dELS) that are >2kb away from transcription start sites, annotated by ENCODE CREs (Moore et al. 2020). We found that proximal-enhancers are under stronger negative selection than distal-enhancers, with a smaller proportion of neutral mutations (76.5% vs. 85.2%, respectively) and a larger proportion of weakly and strongly deleterious mutations (Fig.S2, Table S2c).

While individual mutations in enhancers are inferred to be more neutral than nonsynonymous mutations, the candidate enhancers are enormously large. The active enhancers (EnhA) from ChromHMM as well as enhancers (ELS) from ENCODE each comprise more than 200Mb while non-synonymous sites are only ∼30Mb (Fig. 2a). As a result, the large genomic regions of enhancers can, in aggregate, contribute a larger fitness burden than mutations in coding regions. To investigate this, we calculated the expected number of mutations with specific selection coefficients. We found that although the proportion of weakly deleterious mutations is lower in enhancers compared to non-synonymous mutations, the absolute number of weakly deleterious mutations (10^-5^≤ |*s*|<10^-3^) in enhancers is similar or larger than that for nonsynonymous mutations. Active enhancers from ChromHMM harbor 87% as many weakly deleterious mutations as nonsynonymous sites, while ELS from ENCODE harbor 207% as many (Fig. 2b). We also observed a substantial number of moderately deleterious mutations (10^-3^≤ |*s*|<10^-2^) in enhancers. Active enhancers from ChromHMM harbor 53% as many moderately deleterious mutations as nonsynonymous sites, while ELS from ENCODE harbor 98% as many. These findings suggest that while enhancers have a high prevalence of neutral mutations, their aggregate contribution to deleterious variation is significant, which could have important implications for understanding the evolution of regulatory elements.

### Selection Pressure on Transcribed Regions

We next investigated the DFE for ChromHMM chromatin states association with transcribed regions, including the transcription (Tx), transcription and enhancer (TxEnh), transcription and exons (TxEx), and weak transcription (TxWk) groups of states. We found that the DFEs for Tx, TxEnh, and TxEx were remarkably similar (Fig.S3, Table S2c), with approximately 81.7%, 81.1%, and 79.4% of mutations being neutral, respectively. The weakly transcribed regions (TxWk) had a slightly higher proportion of neutral mutations at 90.9%, indicating reduced selective constraint compared to other groups of states associated with transcribed regions.

The similarity in the DFEs of Tx, TxEnh, and TxEx suggests that regions involved in transcription may experience similar evolutionary pressures, likely due to their shared role in gene expression. In contrast, TxWk regions, which may have lower transcriptional activity, exhibit a larger proportion of neutral mutations, implying less selection. Our results show variable selective pressures across different transcription-related states, reflecting the dynamic and context-dependent nature of transcriptional regulation. Overall, these findings highlight the distinct evolutionary pressures acting on mutations in different categories of noncoding genomic regions.

### Mutations in phylogenetically conserved enhancers and promoters are more deleterious

We next intersected the functional annotations with the PhastCons scores to identify putatively functional noncoding elements that share conserved sequences across primates or mammals.

We first focused on constraint within primates using the PhastCons 17way scores. About 19 Mb of active enhancers annotated by ChromHMM, corresponding to ∼70% of the length of nonsynonymous sites, are highly conserved (top 5% PhastCons scores) (Fig. 2d, Table S2a,f). Within these most conserved enhancers, we infer >50% of new mutations are deleterious (|*s*|>10^-5^). Moreover, in the ChromHMM promoter states within the top 5% PhastCons scores, the proportion of deleterious mutations (|*s*|>10^-5^) is comparable to nonsynonymous mutations, with 75% in conserved promoters versus 76% of nonsynonymous mutations. However, mutations in conserved promoters are still found to be less deleterious overall because most deleterious mutations in promoters are weakly deleterious (10^-5^≤ |*s*|<10^-3^, Fig. 2c, colored in median and dark gray). The proportion of deleterious mutations is smaller in conserved enhancers and promoters when using a less stringent cutoff of the top 10% of the PhastCons scores. However, the difference in the DFE between the top 5% and 10% of conserved sites is small, indicating that deleterious mutations may not be concentrated in the top 5% conserved sites, but are more widely distributed over constraint scores. The least conserved enhancers, corresponding to the bottom 80% of PhastCons constraint scores, are almost neutral, with <=3% of mutations being deleterious (Table S2a). Similarly, for the least conserved promoters, 77% of mutations are neutral (Table S2b).

Next, we focus on more distant phylogenetic conservation in the PhastCons scores from 470 mammalian species. The DFEs of enhancers most conserved in mammals (top 5%) are very similar to the DFEs of enhancers most conserved in primates (Fig.S4, Table S2a). Similar results are also seen for promoters and enhancers annotated by ENCODE (Fig.S4, Table S2a,b). Together, these findings suggest that both functional and comparative genomic annotations provide important complementary information about the fitness effects of mutations.

### Fitness effects of mutations in regions of differing phylogenetic constraint

To better understand the extent to which phylogenetic constraint relates to the fitness effects of mutations in humans, we inferred the DFEs of a broad set of candidate functional noncoding regions across different levels of constraint in primates and mammals. Rather than focusing on individual chromatin states or specific noncoding elements, we examine how phylogenetic constraint relates to negative selection in noncoding regions, which include a mix of candidate functional elements—such as enhancers, promoters, transcribed regions, and other regions with no evidence of being non-functional. We define the candidate functional noncoding regions as noncoding genomic regions excluding ChromHMM quiescent states with weak epigenomic signals and ChromHMM heterochromatin states (Table S1b, Ernst & Kellis 2012; Vu & Ernst 2022). The total length of candidate functional noncoding regions is about 1230Mb. While these bases likely include some non-functional sites, we used a definition that would likely capture many of the known candidate functional regions.

Next, we partitioned the putatively functional noncoding genome into different levels of evolutionary constraint by binning PhastCons scores into 5% intervals (Methods: Binning conserved sites). We then inferred the DFE for each interval (Table S2d). For the candidate functional noncoding regions that fall in the top 5% of the most conserved sites in primates (left side of Fig 3a), we infer that 55% of mutations are deleterious, with the majority (46%/55%) being weakly deleterious (medium and dark gray). We found that the proportion of neutral mutations increases as the phylogenetic constraint in primates decreases (Fig. 3a), corresponding to a decrease in the proportion of deleterious mutations in less conserved regions (Fig.3c, orange line). However, we detect a consistent signal of negative selection in sites in the bottom half of the distribution of constraint scores–sites usually considered to be neutrally evolving (Fig.3a). Specifically, in the bin corresponding of the top 70-75% PhastCons 17way scores, we found that a gamma DFE where 93% of mutations are neutral still fit the data significantly better than a neutral model (Fig.3a, 𝛥𝑙𝑜𝑔 𝑙𝑙 (𝛤, 𝑛𝑒𝑢)). The log likelihood difference is 638, far exceeding our conservative cut-off value of 50 (Methods: DFE inference). A less conserved bin (75-80%), also rejects the neutral model, but contains more than 95% neutral mutations. In even less conserved regions, corresponding to the bins between the top 80% to 85% of primate constraint scores, more than 98% of mutations are neutral. In the final and least conserved bin, a model where mutations were deleterious did not result in a better fit than the neutral model, though the fit of the neutral model also was poor (Fig.S5), suggesting other evolutionary forces at play. We found similar patterns in how DFE changes with the constraint across mammals (Fig.S6c).

**Fig. 3.**
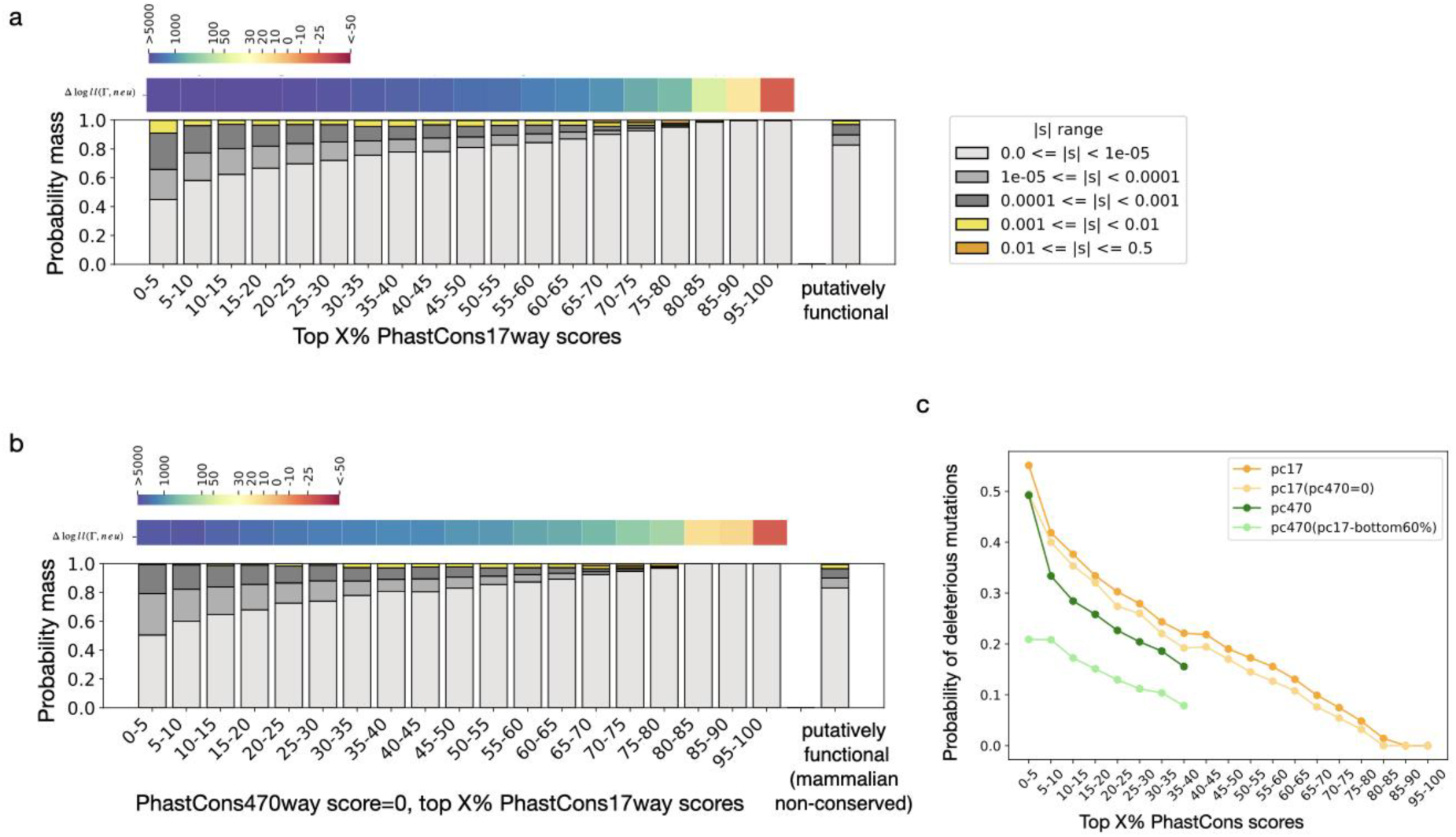
DFEs of candidate functional noncoding regions across different levels of constraint. **a**, The distributions of fitness effects (DFEs) for candidate functional regions across different levels of constraint in primates, measured by PhastCons 17way scores. The left-most bin shows the top 0–5% most conserved regions, and bins become less constrained moving to the right. The 90–95% bin is absent because sites with the smallest non-zero PhastCons scores were grouped into the 85–90% bin, while sites with a score of zero were placed in the 95–100% bin. The far-right bar shows the DFE for the entire curated putatively functional non-coding region (Methods:Processing functional genomic annotations). Vertical bar colors indicate selection strength: light gray for neutral mutations, medium to dark gray for weakly deleterious mutations, yellow for moderately deleterious mutations, and orange for strongly deleterious mutations. The top panel shows the difference in log-likelihood between the gamma model and the neutral model for each corresponding bin. Blue indicates that the gamma model fits the data substantially better than the neutral model, orange indicates comparable fits, and red indicates that the neutral model fits better than the gamma model. **b**, Same as (**a**), but limited to a subset of sites where PhastCons 470way scores equal zero, i.e., sites not conserved across mammals. The last bar showing the DFE for all putatively functional noncoding sites is also limited to the subset of sites with PhastCons 470way scores equal to zero. **c**, Probability of a mutation being deleterious across different levels of evolutionary constraint. As in (a) and (b), the x-axis shows bins of top X% PhastCons scores. The y-axis represents the probability mass of deleterious mutations within each bin, calculated as one minus the proportion of neutral mutations (0<*s*<10^-5^). Orange lines correspond to bins based on PhastCons 17way scores, as in panel (a). The light orange line represents the same 17way bins but restricted to sites with PhastCons 470way scores equal to zero, as in panel (b). The green line shows results for bins defined by PhastCons 470way scores, and the light greenline represents a subset of those sites in the bottom 60% of the PhastCons 17way score distribution.

The proportion of deleterious mutations (defined as 1-the % of mutations in the |*s*|<10^-5^ bin of the DFE) in each bin of constraint scores shows a negative relationship with the constraint scores (Fig.3c). This result suggests that highly constrained genomic regions across primates or mammals are enriched for deleterious mutations, confirming that the phylogenetic constraint can prioritize deleterious mutations. However, this plot shows that even for constraint scores bins in the middle of the distribution (e.g. the top 45-50% PhastCons17way scores), around 20% of the mutations are still inferred to be non-neutral.

Lastly, we estimated the amount of the noncoding genome under selection in humans. We considered mutations with |*s*| >10^-5^ to be deleterious. Overall, within putatively functional regions, we infer 16.9-17.3% of mutations are deleterious (Fig.S6c, 4a, Table S2e). As the putatively functional sites comprise 53% of the human genome, we estimate 9.0-9.2% of mutations in the noncoding human genome are under negative selection. Correcting for a potential upward bias in the inference of the proportion of deleterious mutations (see below and Methods), the estimate drops slightly to 7.7%. The majority (79-85%) of these deleterious mutations are only weakly deleterious (10^-5^<|*s*|<10^-3^).

### Phylogenetic constraint has limited power to identify deleterious mutations

To further understand whether deleterious mutations in putatively functional noncoding regions can be identified by phylogenetic constraint, we examined the cumulative distribution of deleterious mutations as a function of phylogenetic constraint (Fig.4a). Overall, the 5% of the genome that is most conserved across primates carries ∼17.4% of the deleterious mutations. While this represents a clear enrichment, only focusing on the most constrained sites will result in missing the majority of deleterious mutations. Indeed, the top 20% of conserved sites in primates only include 45.5% of non-neutral mutations, the top 30% only include 58.7% of non-neutral mutations, and the top 50% of conserved sites only include about 80% of non-neutral mutations. Using a more strict definition of deleterious mutations (|*s*| ≥10^-4^) returns very similar results (Fig.4a, Table S2h). Similarly, for sites conserved across mammals, the top 5% conserved sites only include ∼14.2% of the deleterious mutations in humans, and the top 40% most conserved sites include ∼52.0% of the deleterious mutations (Fig.S7b, green lines, Table S2h). Similar results were found for enhancers (either annotated by ChromHMM or ENCODE) of which the top 20% of conserved sites in primates or in mammals include less than 60% of the deleterious mutations. For promoters annotated by ChromHMM, the top 20% conserved sites in primates or in mammals include fewer than 50% of the deleterious mutations. However, for promoters annotated by ENCODE, the top 20% of conserved sites in primates or in mammals include 83% or 100% of the deleterious mutations in promoters, respectively (Table S2b). This result likely stems from not identifying many weakly deleterious mutations in the ENCODE enhancers. As the sequence length of promoters annotated by ENCODE is short, we also used a more relaxed cutoff for model selection. The more relaxed cut-off value (𝛥𝑙𝑜𝑔 𝑙𝑙=3, Methods: DFE inference), resulted in more deleterious mutations being identified in ENCODE annotated promoters. Now, the top 20% of conserved sites in primates only include 47% of the deleterious mutations in promoters but the top 20% of conserved sites in mammals include 84% of the deleterious mutations (Table S2b). However, further breaking down the limited sequences of promoters annotated by ENCODE by the level of constraint may result in lack of data to accurately estimate DFEs (Table S2i).

**Fig. 4:**
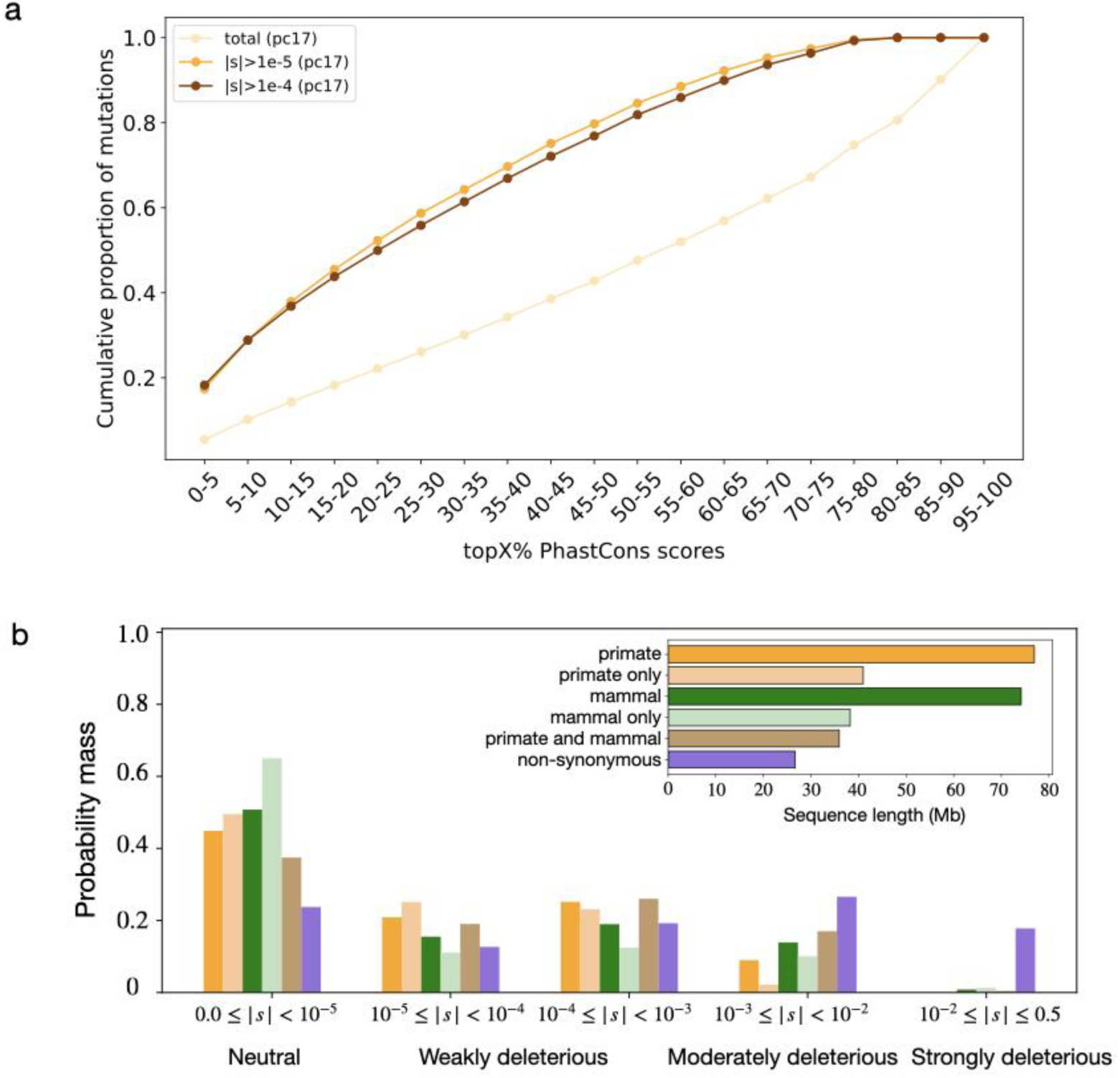
The phylogenetically most constrained regions do not contain the majority of deleterious mutations. **a**, Cumulative proportion of deleterious mutations as a function of primate conservation (17way PhastCons scores). The orange line represents the cumulative proportion of non-neutral mutations across bins. Because the PhastCons scores are discrete, each bin does not contain exactly 5% of the sites. Thus, the cumulative proportion of all mutations is shown for comparison (dark orange). **b**, DFEs of the top 5% most conserved genomic regions. Orange bars represent sites in the top 5% of primate conservation (PhastCons 17way) and pastel orange bars represent a subset of these sites that are not in the top 5% of mammalian conservation (PhastCons 470way). Green bars represent sites in the top 5% of mammalian conservation, with pastel green bars showing those not conserved in primates. Brown bars indicate sites conserved (top 5%) in both primates and mammals. Purple bars show the DFE of nonsynonymous mutations.

Overall, bases conserved across deep evolutionary time are enriched for non-neutral mutations in humans. However, the majority of noncoding deleterious mutations in the human genome are not found within these most constrained regions (top 5%). In sum, our results reveal the limited power of comparative genomic approaches to identify the majority of noncoding deleterious mutations.

### Evidence for changing selective effects over recent time

One potential explanation for the inability of comparative genomic approaches to detect the majority of noncoding deleterious mutations could be changing selective effects over time. Mutations at certain sites may be deleterious in some lineages, but not others, potentially leading to sites not appearing constrained across the phylogeny.

To further investigate this possibility, we examined the DFE of mutations in humans at sites showing conflicting constraint scores between mammals and primates. Sites in the bottom 60% distribution of PhastCons470way scores have a PhastCons 470way score of zero. A zero PhastCons score suggests these sites are not conserved across mammals, and mutations in these sites would often be interpreted to be mostly neutral. However, some sites with a PhastCons 470way score of zero are conserved across primates. For example, across the whole genome, about 20% of the top 5% most conserved sites in primates have a PhastCons 470way score of zero. We also observed a broad distribution of 17way PhastCons scores for sites with a 470way score of 0 (Fig. S7a).

To better understand the fitness effects of mutations in these regions that appear to be constrained in primates, but not in mammals, we inferred DFEs for these genomic regions. Compared to our analyses described above where we only considered a single set of PhastCons scores at a time, we found more neutral mutations in genomic regions not conserved across mammals, even though they have the same level of constraint in primates (Fig. 3a,b). Nonetheless, despite the lack of constraint across mammals, ∼50% of mutations in sites highly conserved across primates (top 5% PhastCons17way scores) were deleterious in humans. We also found evidence of negative selection at mutations in sites ranked up to the top 70%-75% conserved across primates (Table S2d). In fact, mammalian non-conserved sites exhibited only a slightly lower proportion of deleterious mutations compared to all sites with similar primate constraint (Fig. 3c, orange versus light orange line). These results suggest that a proportion of mutations at sites conserved across primates, but not conserved across mammals, are still deleterious.

As a control, we performed the reciprocal analysis, where we inferred the DFE for different bins of mammalian constraint, focused on sites not conserved in primates. Here, we define sites not conserved in primates to be those in the lower 60% bins of the distribution of primate constraint scores which yields a similar proportion of sites as those not conserved in mammals (i.e. PhastCons 470way score equals zero). Inferring the DFE for mutations in the 5% most constrained sites in mammals that are not constrained in primates reveals only 20% of mutations to be deleterious (Fig. 3c light green line, Fig. 6d). This is substantially smaller than the proportion of deleterious mutations inferred (50%) for the 5% most constrained sites in mammals, without regard to primate constraint (Fig. 3c dark green line, Fig. 6c). More broadly across the distribution of mammalian constraint, the proportion of deleterious mutations (|*s*| >10^-5^) is substantially lower for sites constrained in mammals, but not constrained across primates (Fig.3c, two green lines). These results suggest that mutations in noncoding genomic regions not conserved across mammals can still be deleterious in humans. Thus, the primate PhastCons scores may better reflect constraint in the human population than the mammalian PhastCons scores.

The fact that mutations in regions of the genome showing little cross-species constraint are inferred to be deleterious suggests that there may be changing selective pressures over evolutionary time. This hypothesis predicts that mutations at sites constrained across primates ought to be under more selection in humans than sites that appear constrained across mammals as a whole. To test this, we stratified bases by PhastCons score into 5 categories (Fig. 4b) and inferred the DFE for each category. Overall, mutations in regions that are constrained in both mammals and primates are more deleterious than mutations constrained in only one set of species. However, sites only conserved in primates (orange) are more likely to be deleterious than sites that are only conserved in mammals (green). However, the proportion of moderately (0.001≤ |*s*|<0.01) and strongly deleterious (0.01≤ |*s*|<0.5) mutations is greater for sites constrained across mammals compared to primates (Fig. 4b, Table S2g). These findings suggest that some functional noncoding sites are experiencing natural selection in primates, but not the entire mammalian phylogeny, supporting the hypothesis of turnover of selection across evolutionary time.

### Validation of the DFE estimation

To validate our inferences of the DFE and assess robustness to complications like complex genome structure and linked selection, we conducted two additional studies. First, we performed bootstrapping to understand the uncertainty of our DFE estimates. Second, we carried out forward in time simulations under different scenarios and then inferred the DFE using the same approach as we applied to the empirical data.

We first assessed the uncertainty of our estimate of the DFE, especially for genomic regions with a large proportion of neutral mutations. Here we resampled 100kb regions of the genome and inferred the DFE of the ChromHMM active enhancers on these bootstrapped datasets (see Methods: Bootstrapped Enhancers). We focused on the ChromHMM active enhancer regions as we inferred there to be a relatively high proportion (2.5%) of strongly deleterious mutations (Fig. 2a). We found the DFEs of bootstrapped enhancers are nearly the same as for the original ChromHMM active enhancers (Fig.S9), suggesting little statistical uncertainty in our parameter estimates. This result indicates that the 2.5% of mutations in ChromHMM active enhancers inferred to be strongly deleterious is not due to statistical uncertainty due to a limited number of sites.

Second, we assessed performance of our DFE inferences on simulated data with a known DFE. We simulated data with a human genome annotation including coding regions, conserved noncoding regions, quiescent sites, and the rest of the genome as background regions (see Methods: Simulations to assess performance of the inference). Each simulation replicate contained ∼130 Mb of background regions and 9Mb of conserved noncoding regions. We examined various DFEs for the background and conserved noncoding regions, while the DFE of non-synonymous mutations is from Kim et al. 2017, and mutations at the quiescent sites are always neutral.

We simulated a selection model in which conserved noncoding regions follow the inferred DFE from the top 5% most conserved sites in mammals (Table S3b,d, where 50% of mutations are neutral), while background regions follow a DFE where 90% of mutations are neutral (Table S3a,c). In all simulation replicates, we confidently distinguished between the neutral model and the selection model, even when the proportion of neutral mutations is 90% (Table S3e). Also, we accurately estimated the proportion of neutral mutations, with a 4% underestimation when comparing the mean inference of simulated replicates with the simulated DFE of background regions (Fig. S8a, Table S3a). We slightly overestimated the proportions of deleterious mutations at all the selection coefficients for background regions, however. The deviation ranges between 0.4% for strongly deleterious mutations (|*s*|>0.01) to 1.7% for 10^−5^≤ |*s*|<10^−4^). For conserved noncoding regions, we also slightly underestimate the proportion of neutral mutations (−2.0%) and weakly deleterious mutations (−0.40%), while overestimating moderately and strongly deleterious mutations (+1.6% and +0.75%) (Fig. S8b, Table S3b). We also observed a stronger underestimation of neutral mutations and overestimation of deleterious mutations in both background regions and conserved noncoding regions when simulations were performed with a mutation rate twice as high as the one used here (Fig. S8c,d; Table S3c,d). This result indicates that the complicated linked selection in the human genome might bias the estimates.

To test whether our conclusions are robust to potential bias in DFE estimates, we scaled DFEs of enhancers, promoters, and candidate functional noncoding regions binned by PhastCons scores according to the bias found in simulated data (see Methods: Scale DFE based on simulations). We found that the DFEs of enhancers and promoters (Fig. S10a, Table S3f) are largely unchanged, and the number of weakly deleterious mutations in enhancers is comparable to the number in non-synonymous mutations (Fig. S10b, Table S3f). Similarly, we still observe a decrease in deleterious mutations in less conserved sites and a large drop when sites are not conserved across primates (Fig. 3c and Fig. S10c). In addition, phylogenetic constraint still has limited power to identify deleterious mutations (Fig. S10d, Table S3i). For example, the top 20% of conserved sites in primates only include 50% (45.5% before correction) of non-neutral mutations. Thus, our conclusions are robust to potential bias in estimating DFEs.

## DISCUSSION

Here we leveraged recent functional genomic annotations and estimated the fitness effects of mutations in enhancers, promoters, and conserved noncoding regions. We found that while most mutations in enhancers are often neutral, approximately 30% of new mutations in promoters are deleterious. We then intersected the functional genomic annotations with patterns of phylogenetic constraint across mammals and primates. Intriguingly, the most evolutionarily conserved sites across mammals—frequently used to pinpoint functional regions in genomic studies—encompass only a small fraction of deleterious mutations. For instance, the 5% most conserved noncoding sites in mammals contain less than 20% of deleterious mutations. This suggests that weakly deleterious genetic variation is more widely distributed across functional noncoding genomic sites than previously appreciated (Mouse Genome Sequencing Consortium et al. 2002; Siepel et al. 2005; Lindblad-Toh et al. 2011; Arbiza et al. 2013; Gulko et al. 2015; Dukler et al. 2022). Additionally, we observed evidence of selection coefficients changing over deep evolutionary time, suggesting a dynamic architecture of gene regulation.

Our work builds on previous work studying the fitness effects of noncoding mutations. While numerous studies using human polymorphism data have detected the effects of negative selection on noncoding mutations (Keightley et al. 2005; Drake et al. 2006; Asthana et al. 2007; Katzman et al. 2007; Eory et al. 2010; The ENCODE Project Consortium 2012; Arbiza et al. 2013), most of these studies did not estimate a full DFE of deleterious mutations. Not having a full DFE limits quantitative comparisons regarding the extent of selection in different annotations. Further, a DFE is necessary for future population genetic models and simulations of noncoding variation. One study, however, inferred the DFE for human-mouse conserved noncoding sequences flanking genes. They inferred a DFE where approximately 76% of mutations were neutral (|*s*|<10^-5^), ∼8% of mutations were in each of the 3 weakly to moderately deleterious bins, and 1.5% of mutations were strongly deleterious (|*s*|>0.01). These proportions align well with what we found in our analysis using functional genomic annotations (Fig. 2a). For example, for enhancers, we inferred approximately 80-90% of mutations to be neutral while for promoters, ∼70% of mutations were inferred to be neutral. While enhancers comprise a greater number of basepairs, we still would still expect the DFE inferred by Torgerson et al. for human-mouse conserved regions to be more deleterious than we infer for enhancers as the former focused on conserved cis-regulatory elements. Our study moves beyond Torgerson et al. (2009), by studying noncoding variation across the genome, rather than focusing exclusively on human-mouse conserved regions, like Torgerson et al. did. Additionally, we leverage modern functional genomic annotations of putative enhancers and promoters to increase the resolution of our analysis.

Comparative genomic studies are widely used to identify noncoding sites where mutations are deleterious. Our results indicate that highly conserved sites across multiple species (i.e. the top 5% most conserved sites in mammals and primates) are enriched with deleterious mutations (Fig 3). For example, we infer 15% of mutations in the 5% most conserved sites across 470 mammals are moderately to strongly deleterious (*s*<-0.001, Table S2d). For sites that are less conserved, the DFE is skewed to be more neutral. However, even considering up to ∼⅔ of the genome, we find that a model with deleterious mutations fits substantially better than a fully neutral model, suggesting widespread prevalence of weakly deleterious mutations, even outside of the most conserved sites. Viewed another way, fewer than 20% (Table S2d) of the deleterious noncoding mutations in humans occur in the 5% most conserved sequence.

These findings suggest a fundamental limitation to comparative genomic approaches across broad phylogenetic scopes. Mutations at some sites may be subject to relaxed selection in certain lineages. This may erode the signal of constraint when considering a large phylogeny (e.g. 470 mammals). Thus, mutations at sites that are not conserved at a broader phylogenetic level may still be under negative selection in specific lineages. Conversely, sites that are conserved across a broad phylogeny may not continue to experience selection in all clades.

Interestingly, and consistent with the prediction outlined above, we observe that sites highly conserved across primates exhibit a greater proportion of deleterious mutations in humans compared to those sites conserved only across mammals but not specifically highly conserved in primates. Thus, if the aim is to identify noncoding mutations that are more likely to be deleterious in humans, focusing on conservation within more closely related species might be more effective for capturing the largest number of deleterious mutations. We note that further work is needed to probe the relationship between fitness effects in humans and phylogenetic constraint across species as multiple factors could be at play. For example, the phastCons 17way scores and the phastCons 470 way scores are based on alignments generated by different alignment algorithms (MultiZ vs. Cactus) which is a potential confounder when making primate vs. mammal differences particularly related to issues of alignment noise.

Moreover, we found that highly conserved sites in primates are more likely to harbor weakly deleterious mutations, which could potentially serve as causal variants for complex diseases. Conversely, mutations at sites conserved across mammals are more likely to be moderately to strongly deleterious. Depending on the objective—whether identifying weakly or strongly deleterious mutations—the phylogenetic context of conservation should be considered. Including more species is not necessarily better; it is important to consider the phylogenetic scope.

The limitations of comparative genomic approaches suggest the importance of polymorphism-based constraint scores (like the Broad Gnochi scores and DR scores, Chen et al. 2024; Halldorsson et al. 2022) to identify deleterious mutations in humans. These scores leverage large samples of human genomes (>10,000 individuals) to detect regions of the genome with a depletion of polymorphism as being subject to negative selection. Such studies have the advantage of measuring constraint directly in the human lineage, rather than relying on distantly related lineages, as comparative genomic approaches do. However, these scores generally are at a kb resolution, which is much lower than comparative genomic scores based on divergence. As such, there are still challenges to identify individual noncoding deleterious mutations in the human genome.

Our results have a number of implications for population genetic studies of genetic variation. First, inferences of demographic history typically require genetic variants that are neutrally evolving. To find such sites, researchers may focus on noncoding regions that are not in the most phylogenetically conserved bases (Gronau et al. 2011; McManus et al. 2015; King & Wakeley 2016; Veeramah et al. 2018). Our finding that variants outside of the most conserved bases may be deleterious implies that filtering based on constraint may miss variants under selection. Instead, we suggest more aggressive filtering, such as removing >20% of the phylogenetically constrained bases and bases not in quiescent or heterochromatin states in ChromHMM. These filters could be used in addition to previously proposed filters for background selection and biased gene conversion to obtain the most neutral set of variants possible. Second, there has been interest in assessing how recent human demography affects patterns of deleterious variation across populations (Lohmueller 2014; Simons et al. 2014; Henn et al. 2015; Peischl & Excoffier 2015; Henn et al. 2016; Simons & Sella 2016; Koch & Novembre 2017; Kyriazis & Lohmueller 2024). To date, these studies have primarily focused on nonsynonymous variants. Our finding of a significant accumulation of weakly and moderately deleterious variants in noncoding regions suggests that noncoding sites might contain an important fraction of the genetic load. As noncoding mutations are more nearly neutral compared to nonsynonymous variants, the former may be more affected by different amounts of drift in different populations. Future work could use our inferred DFEs for non-coding mutations in simulation studies to investigate the impact of differences in demography on genetic load.

We attempted to control for various confounding factors that could affect inferences of the DFE from genetic variation data. First, background selection can be a pernicious factor that affects patterns of variation. Deleterious mutations in coding regions and conserved noncoding elements may impact the nearby linked neutral variation. In principle, this could confound our inferences of the DFE on noncoding non-conserved sequences. In other words, could the effects of negative selection that we infer for non-conserved noncoding sequences actually be the result of background selection on conserved noncoding or coding sequences? Several lines of evidence suggest that this is not the case. First, the demographic model that we infer from putatively neutral sites provides some degree of correction for background selection (Kim et al. 2017). We developed a novel approach (see Methods) to pick nearby flanking quiescent sites to use to infer a demographic model. This demographic model is then used in the DFE inferences. Because these quiescent sites are located nearby the functional (and putatively selected) sites, they should experience a similar amount of background selection. Consequently, the demographic models we infer will account for this background selection and the DFE we fit to the functional sites will capture the residual signal due to the direct effects of selection. Second, we have conducted forward in time simulations including linkage to assess the impact of background selection on the inference. We found that with realistic human mutation rates, there is little bias due to background selection. With elevated mutation rates (2-fold larger than the realistic human mutation rate), background selection appears to have more of an effect. Importantly, we found that the gamma model was still preferred over a neutral model for up to the top 80% of conserved sites across primates, even when using the maximum likelihood cutoff obtained from our high-mutation-rate neutral simulation as the cutoff for model selection. Thus, background selection alone should not lead to this strong of an improvement in fit to the selection model over the neutral model. As such, our results are not attributable to background selection from linked deleterious mutations at conserved noncoding or coding sites.

A second confounder is mutation rate variation along the genome, particularly in different functional annotations. Indeed, it is likely that mutation rates covary along the genome with different chromatin states (Seplyarskiy et al. 2022). Our inferences of the DFE use the numbers of SNPs at different frequencies in the sample. As such, an accurate mutation rate is important for reliable inferences. Thus, we accounted for mutation rate variation by using a high-resolution mutation rate map (Seplyarskiy et al. 2022). While we cannot rule out more subtle effects escaping this correction, our approach to model mutation rate variation along the genome should capture the most important variability in polymorphism attributable to mutation rate variation. As such, remaining variation in levels of polymorphism across the genome can be attributed to direct and indirect effects of natural selection.

Since the completion of the mouse genome project in 2000 (Mouse Genome Sequencing Consortium et al. 2002), researchers have been interested in estimating the proportion of bases in the human genome where mutations are deleterious. Initial estimates from comparative genomic studies were around 5-6% (Mouse Genome Sequencing Consortium et al. 2002; Cooper et al. 2005; Siepel et al. 2005; Pollard et al. 2010; Lindblad-Toh et al. 2011), with a later one increasing to 7% (Davydov et al. 2010). Subsequent studies, using a variety of patterns in the data and models, have suggested potential functional turnover, leading to mutations at certain positions being deleterious in some lineages but not in others. Models accounting for turnover suggested a greater proportion of bases where mutations were deleterious, with estimates >=8% (Meader et al. 2010; Ponting & Hardison 2011; Ponting et al. 2011; Rands et al. 2014). Taking a different approach, Ward and Kellis (Ward & Kellis 2012) leveraged early polymorphism datasets from the 1000 Genomes Project and the functional annotations from the ENCODE project and inferred negative selection acting on mutations at functional sites that were not phylogenetically conserved across species. In total, they estimated that ∼9% of the human genome is under negative selection, with nearly half being outside of phylogenetically constrained regions. While this study received some criticism due to technical artifacts in the 1000 Genomes data as well as not fully accounting for background selection (Green & Ewing 2013), the overall conclusions of Ward and Kellis are in line with ours. Namely, we infer that 9.0%-9.2% (Table S2e) of noncoding bases in the human genome contain mutations that are deleterious in humans. While this finding was controversial in 2012, subsequent comparative genomic studies using 240 species (Christmas et al. 2023) also revised the amount of the human genome subject to negative selection. They inferred that ∼10.7% (332 Mb) of the bases in the human genome are constrained by comparing the distribution of constraint scores across the human genome to those seen in neutral ancestral repeat sequences. Curiously, their phylogenetic approach to identify constrained elements was only above to recover ∼30% of the constrained bases (101Mb in constrained elements at FDR of 0.05 out of a total of 332 Mb of constrained sequence). Thus, these two findings together imply that comparative genomics alone cannot locate the majority of the bases under selection in the human genome. Considering the totality of the evidence, it is time to revise the estimate of the amount of the human genome that is under selection. The best available evidence suggests that the true proportion of the genome under selection is closer to 9%, almost 2-fold higher than the early estimates of 5%. New datasets and models will be required to discover these positions containing deleterious mutations and determining their contribution to evolution and phenotype.

## METHODS

### Processing functional genomic annotations

For all the functional genomic annotations, either from ChromHMM or ENCODE, we removed all the coding regions annotated by GENCODE (Frankish et al. 2022). We used a previously defined universal ChromHMM chromatin state annotation of the human genome (hg38 liftOver) (Vu & Ernst 2022). This annotation consisted of 100 states which were previously grouped into 16 major groups based on biological characteristics. For the analyses here, we defined the major group EnhA as active enhancers. Promoters were defined as a combination of three major groups: TSS, PromF, and BivProm. We defined the putatively functional noncoding genomic regions as those that do not include sites annotated as being in the quiescent chromatin state (Quie) or heterochromatin state (HET). In addition, in our analyses of putatively functional noncoding genomic regions, we also only kept genomic sites that have both PhastCons 17way and PhastCons 470way scores. We downloaded all the human enhancers (cCRE-ELS), promoters (cCRE-PLS), proximal-enhancers (cCRE-pELS), and distal-enhancers (cCRE-dELS) from 1518 cell types annotated by ENCODE cCRE from SCREEN (Moore et al. 2020). The promoters (cCRE-PLS) are cCREs that fall within 200 bp (center to center) of an annotated GENCODE transcription start site and have high DNase and H3K4me3 signals. The enhancers (cCRE-ELS) are cCREs that have high DNase and H3K27ac signals, but a low H3K4me3 signal if they are within 200 bp of an annotated transcription start site. Proximal-enhancers (cCRE-pELS) are the subset of cCREs-ELS within 2 kb of a transcription start site, while the remaining subset is denoted as distal-enhancers (cCRE-dELS) (Moore et al. 2020).

### Binning conserved sites

We downloaded PhastCons 17way and PhastCons 470way scores from the UCSC Genome Browser. Each site of the 2,656,464,721 bp has a PhastCons 470way score and each site of 2,669,831,920 bp has a PhastCons17way score. We split all the genomic sites with PhastCons 17way scores, including coding sites and sites that we later removed from downstream analyses, into bins, each of which contains approximately 5% of the genomic sites that have PhastCons 17way scores. We separately binned genomic regions with PhastCons 470way scores the same way. However, many sites have the same PhastCons score, and thus we could not split the sites evenly into bins containing exactly 5% of the sites. Thus, we determined an empirical score cutoff for each bin based on a random subset of the sites. We included the score of the lower percentile, for example, the cutoff for the top 5% of the PhastCons17 score is 0.839 and the cutoff for the top 10% is 0.533. Then, the top 5-10% bin included sites with scores that are within [0.533,0.839). The thresholds used in splitting the genome are listed in Table S1d. The genomic regions with PhastCons 17way scores are binned into genomic sites with top 0-5%, 5-10%, 10-15%, … , 80-85%, 85-90% and 95-100% PhastCons 17way scores. We retained 19 bins because the cut-off value for both the bin of 85-90% and 90-95% was 0.001, the smallest non-zero PhastCons 17way score. Thus all the genomic sites with the PhastCons17way score of 0.001 were included in the bin of 85-90%. Thus the bin of 85-90% includes 7.1% of total sites that have PhastCons17way scores. The rest of the bins have between 4.86% to 6.94% of the sites and the mean proportion of sites of all the 19 bins is 5.26% (Table S1e). We then curated the putatively functional noncoding genomic regions (as defined above) with different levels of constraint based on the 5% bins of PhastCons 17way scores. If PhastCons scores are evenly distributed across the genome, we would expect to see the same proportion of putatively functional noncoding sites in each bin. However, the putatively functional noncoding regions are enriched in highly conserved sites. 6.31% of putatively functional genomic sites have the PhastCons scores in the top 5% bins, compared to 5.02% for the whole genome (including coding regions). Each bin of PhastCons17way scores contained 4.44% to 7.7% (average of 5.3%) of the total number of putatively functional noncoding sites (Table S1e).

Similarly, we also binned genomic sites based on PhastCons 470way scores. Because the PhastCons 470way scores had been rounded to three decimal places, about 60% genomic sites have a PhastCons 470way score of zero. Thus, we are only able to split the genome to the top 0-5%, 5-10%, 10-15%, …, 35-40% based on non-zero PhastCons 470way scores. For the whole genome, each bin of non-zero PhastCons470way scores contained 3.64% to 8.02% (average of 5.47%) of the total number of sites. For the putatively functional noncoding sites, proportions ranged between 2.90% to 6.6%, with a mean of 4.82% for PhastCons470way non-zero scores. Similarly, a larger proportion (6.09%) of putatively functional sites are found to have the top 5% PhastCons 470way scores compared to the whole genome (5.00%). We split the remaining 60% of the sites by the 19 bins of top X% PhastCons 17way scores.

### Calculating mutation rates

We calculated the per-site per-generation haploid mutation rate (*μ*) at basepair resolution based on the Roulette mutation rate map (http://genetics.bwh.harvard.edu/downloads/Vova/Roulette/), which provides mutation rates of all three possible single-nucleotide changes at each site (Seplyarskiy et al. 2022). For each genomic position, we summed up the three mutation rate (“MR”) values and scaled the total by 1.015×10^-7^ following the authors’ recommendation (https://github.com/vseplyarskiy/Roulette/tree/main/adding_mutation_rate), to obtain the diploid mutation rate. We then divided the diploid rate by two to derive the haploid per-site per-generation mutation rate. Across our test groups, the estimated mutation rates (*μ*) varied by approximately two-fold, ranging from ∼9×10^-9^ to ∼2×10^-8^.

### Analyzing genetic variation

We used 108 Yoruba (YRI) individuals and downloaded the variant call files from the 30x coverage NYGC 3202 samples of the 1000 Genomes Project sample collection (1000 Genomes Project Consortium et al. 2015; Byrska-Bishop et al. 2022). We retained only biallelic SNPs located on autosomes within high-quality noncoding regions, as defined by the genome accessibility mask from the 1000 Genomes Project.

We generated the site frequency spectrum (SFS) and calculated polymorphism patterns using functions implemented in *dadi*. To account for the possibility of missing data, we projected the SFS from the total of 216 chromosomes (108 YRI individuals) at each site to a sample of 160 chromosomes using dadi.Spectrum.from_data_dict, projection=108. Sites with data from less than 160 chromosomes were removed. We calculated per-site heterozygosity *pi*, and Tajima’s *D* by functions implemented in *dadi* (fs.pi,fs.Tajima_D()). However, the sequence length and mutation rates vary between functional annotations. Thus, we calculated *pi_norm* by dividing *pi* by *μ*𝐿, where *μ* is the haploid per-site per generation mutation rate obtained from the Roulette mutation rate map (see the above), and 𝐿 is the number of annotated sites.

### DFE inference

We inferred the DFE from the SFS using a maximum likelihood approach with two steps. First, we inferred the demography based on SFS from quiescent control sites which are presumably more likely to be neutrally evolving than annotated functional sites. Second, we inferred the parameters of the DFE of the test set, which included various putatively functional noncoding mutations we defined in the Methods, conditional on the demographic inference. This two-step approach can effectively control for the effects of demography and linked selection when estimating the DFE of new mutations (Kim et al. 2017). We implemented both the demographic and DFE inferences in dadi-cli (https://github.com/xin-huang/dadi-cli) (Huang et al. 2023)

### Models of demography and the DFE

We fit a three-epoch model with two population size changes to the quiescent control sites. This model included four parameters: the ratio of the bottleneck to the ancestral population size (*nuB*), the ratio of the contemporary to the ancestral size (*nuF*), the duration of the bottleneck (*TB*), and the time since recovery (*TF*) (Fig.S11a). *nuB* and *nuF* are measured in units of the ancestral population size, Nₐ, and *TB* and *TF* are measured in units of 2Nₐ generations. *nuB* and *nuF* can take values greater than or less than one.

For the test sites, we estimated the distribution of population-scaled heterozygote selection coefficient (2Nₐ*s*). We assumed several different functional forms of the DFE including a neutral, a gamma, and a neugamma distribution. In the neutral model, all the mutations are neutral with a selection coefficient (*s*) equal to 0. The gamma distribution has a shape parameter and a scale parameter. The neugamma distribution is a mixture distribution consisting of a point mass of neutral mutations with *s*=0 plus a gamma distribution. We estimated the parameters of these parametric distributions but, for ease of visualization, present a 5-bin discrete DFE in the results.

### Inference of demography using quiescent sites

We used sites annotated as quiescent by ChromHMM to infer demographic history. Although not all mutations at quiescent sites are entirely neutral, mutations at these sites as a group are more likely to be neutrally evolving than mutations at sites annotated in a functional state. To increase the likelihood that we only primarily include neutrally evolving mutations, we excluded coding regions and sites annotated as human enhancers or promoters by ENCODE, retaining only noncoding quiescent sites as the control regions for demographic inference. To control for the effects of linked selection and broad-scale mutation rate variation, we selected quiescent sites that were nearby our test sites. To do this, for each contiguous region of test functional sites, we selected all the nearby quiescent sites within a 100 kb window centered on the middle of the test sites. We combined all the nearby quiescent sites of all the continuous regions of test sites, only retaining each quiescent site once if it was chosen multiple times. These control sites are expected to share similar genomic features, such as mutation rate, recombination rate, gene density, and linked selection, with the test sites. The window is large enough that most test sites had suitable controls. We excluded test/functional regions without nearby control sites, but we still kept at least 90% of the test sites in all genomic regions analyzed.

### Modeling mutation rate variation

Inferences of the DFE require an estimate of the population scaled mutation rate, *θ*. This quantity is related to the expected number of mutations in the test sites. Because of the differing sequence lengths and mutation rates between the test and control groups, the expected number of mutations may differ between the test and control groups. Therefore, we adjusted the expected SFS of the test group by multiplying it by the ratio of mutation numbers between the test and quiescent control groups. We also scaled the population-scaled selection coefficient (Nₐ*s*) to the selection coefficient (*s*) by Nₐ estimated from *θ* of quiescent regions, using the equation *θ* = 4Nₐ*μ*L, where *μ* is the average mutation rate and *L* is the sequence length of the quiescent control regions. The value of *θ* was directly inferred using *dadi*.

### Finding the maximum likelihood parameters of demography and selection

We used *dadi* to infer demographic history and Fit*dadi* to infer the DFE from the SFS. Both methods perform numerical optimization of the likelihood function and require a set of initial parameter values with optional user-defined parameter ranges. *dadi* then performs parameter perturbations followed by optimization to identify the best-fit model. While the final result should, in theory, be independent of the initial parameters, we found that the choice of starting values could influence the outcome. In particular, some initializations failed to recover the global maximum likelihood. To address this, we performed inference using a grid of initial parameters spanning the specified ranges.

For fitting the three-epoch demographic model to the empirical data, we used parameter bounds of 1 to 10 for *nuB*, 2 to 1000 for *nuF*, and 10^-7^ to 2.0 for both *TB* and *TF*. Based on initial runs, we restricted the range for generating initial values to be focused toward the regions of the parameter space that yielded robust fits. Specifically, we used a narrower range of 1.0 to 2.0 for *nuB*, 0.5 to 0.9 for *TB*, and 10^-7^ to 0.1 for *TF*. We generated four linearly spaced values within each range, resulting in 256 combinations of initial parameters. For fitting simulated data with the two-epoch model, we used parameter bounds of 0.5 to 5 for *nu* and 10^-5^ to 10^5^ for *T*. We generated initial values within these ranges using eight logarithmically spaced points for each parameter, excluding the boundary values.

For DFE inference, we used the same initial parameters in both empirical analyses and simulations. For the gamma distribution, we generated 30 log-spaced shape parameters ranging from 10^-10^ to 2, and 30 log-spaced scale parameters ranging from 10^-10^ to 10^8^. This parameter range covers a broad range of DFEs, especially DFEs with a large proportion of neutral mutations (Fig.S11). To avoid unnecessary calculations, we retained only combinations whose mean population selection coefficients, calculated as the product of shape and scale, fell within a biologically plausible range of 2Nₐ*s* between 2Nₐ*s*×10^-5^ and 2Nₐ*s*×0.5, corresponding to approximately 0 to lethality.

For the neugamma distribution, which includes additional parameters, we generated 10 log-spaced shape parameters (10^-10^ to 2), 10 log-spaced scale parameters (10^-10^ to 10^8^), and 10 linearly spaced values for *p_neu_*, the proportion of neutral mutations, ranging from 0 to 1. Similarly, we retained only shape-scale combinations whose mean selection coefficient fell within the defined biologically plausible range.

To easily run the inference in high-throughput manner, we used the command-line package *dadi*-cli (Huang et al. 2023) based on the version of *dadi* that includes the Fit*dadi* module for DFE inference. However, the neugamma model has not been specified, so we modified the packages to fit neugamma model. The modification can be found in github (https://github.com/chenludi/DFE_noncoding/tree/main/modify_dadi_dadi-cli).

### Selecting the best-fit DFE model

We selected the best-fit DFE model for each annotation based on comparison of the log-likelihoods of the neutral, gamma or neugamma models. The cut-off value of log-likelihood difference for selecting the best-fit model can be found through a likelihood-ratio test (LRT). The likelihood ratio, LR=2(log_ll_model1-log_ll_model2), where log_ll_model is the log likelihood value of the model. Asymptotically, the LR follows a χ² distribution under the null hypothesis. Thus, the p-value can be calculated by 1 - chi2.cdf(LR, df) using the chi2 function from the python package scipy.stats. A 5% significance level (p < 0.05) corresponds to a log-likelihood difference of 3 when comparing the gamma and neutral models (df=2) and corresponds to a log-likelihood difference of 2 when comparing the gamma and neugamma models (df=1). However, the asymptotic distribution of the LRT under the null assumes that the SNPs are independent of each other. The linkage between SNPs violates the independence required for a χ²-based likelihood-ratio test. In addition, long sequences with more data risk over-fitting gamma or neugamma models. Thus, we needed a more strict threshold for selecting the best-fit model.

Our forward simulations where all noncoding mutations are neutrally evolving (see below) allowed us to define a conservative cutoff for model selection. In the simulation, the background region is about 130Mb which is close to the length of most test groups in our analyses. For example, each bin of the PhastCons scores has a sequence length between 55-100Mb. The conserved noncoding regions in the simulation were approximately 9Mb, which is similar to some of the test groups with smaller sequence length, such as promoters annotated by ENCODE. In these null simulations, the maximal improvement in log-likelihood when fitting the gamma model over the neutral model was 35 log-likelihood units for the long background regions, with a 95% confidence interval to be [9.0, 21.8] (Tabel S3h). For conserved noncoding regions, the log-likelihood difference is smaller than 0.1. Thus, we selected a critical value of 50 log-likelihood units as a conservative criterion to avoid false detection of negative selection in empirical analyses with a similar length to those used empirically. However, for promoters annotated by ENCODE (PLS) binned by PhastCons scores, we also used a smaller cut-off value of 3 (5% significance level, df=2) for comparison because the sequences are smaller than 1.5Mbp.

When considering a higher mutation rate of 2×10^-8^, the effects of linkage were more pronounced (Fig.S8c,d, Table S3c,d). When comparing the fit of the gamma model to the neutral model, we observed a maximum improvement in log-likelihood of 262 for background regions and 15 for conserved noncoding regions (Table S3h). These results indicate a higher risk of overfitting with more segregating variants. Importantly, in the model comparisons on the empirical data, we observed large (>450 log-likelihood units) improvements in fit for the gamma model over the neutral model for all but the lowest (>80%) phastCons bins, suggesting that linkage cannot account for the improvement in fit of models with selection.

### Bias-corrected DFEs based on simulations

As we found a slight upward bias in the proportion of deleterious mutations when analyzing simulated data (Fig. S8), we re-scaled our inferred DFEs to remove this bias. Specifically, we scaled the 5-bin discrete DFEs of enhancers, promoters, and candidate functional noncoding regions binned by PhastCons scores according to the bias found in simulated data. To do this, we first calculated the ratio of the true proportion of mutations at a given range of selection coefficient, e.g. 0≤ |*s*|<10^-5^, and the mean estimates from bootstrap replicates. For simulated background regions with 90% neutral mutations, the ratios between true values and estimates of bootstrap replicates are 1.05, 0.65, 0.78, 0.77, 0.30 for five bins of selection coefficients (Table S3f). A ratio larger than 1 means the true value is larger than the mean of the inferred values. We scaled the empirically inferred DFEs with more than 75% neutral mutation using this ratio. For annotations where we estimated fewer than 50-75% of the mutations to be neutral, we scaled the DFE by the ratios from simulated conserved noncoding regions, which have a similar proportion of neutral mutations. Here, the ratios between true values and estimates of bootstrap replicates are 1.04, 1.02, 1.00, 0.90, 0.53. In the simulations and most inferred DFEs using functional annotations, the proportion of mutations whose selection coefficient is larger than 0.5 is almost zero. For the few that are not zero, we added the proportion to the strongly deleterious with 0.01 ≤ |*s*| ≤ 0.5. After scaling the estimates by the ratio, we normalize proportions so that proportions sum to 1.

### Bootstrapped Enhancers

To quantify the variance in allele frequency distributions across enhancer regions, we implemented a genome shuffling approach. First, the entire genome was segmented into contiguous 100-kilobase blocks. Next, a bootstrapping procedure was applied—sampling these blocks with replacement (allowing for duplicated segments)—to construct a randomized genome. Finally, the actual genomic coordinates of enhancers were mapped onto this bootstrapped genome, yielding a randomized set of enhancer regions for subsequent analysis.

### Estimate the average DFE functional noncoding regions

The average DFE of the entire putatively functional noncoding region was calculated as a weighted average of DFEs across genomic regions binned by their PhastCons 17way scores (Fig. 3a) or binned by both PhastCons 17way scores and PhastCons470way scores (Fig. S6c, Fig. 3b). For each bin, the DFE was weighted by its estimated mutation number, calculated as the product of the average mutation rate and the sequence length of that bin.

The proportion of non-neutral mutations for the whole human noncoding genome was calculated as a weighted average of putatively functional genomic regions and the rest noncoding genomic regions. Instead of weighted by mutation number, we weighted by sequence lengths. The putatively functional genome is about 1.230×10^9^ bp and we consider the whole noncoding genomic region to be 2.3× *10^9^*. Thus, the proportion of the genome that is under selection is the product of the proportion of non-neutral mutations (∼0.17) for the whole genome times the sequence length in putatively functional noncoding regions divided by the sequence length of the whole noncoding genome.

### Simulations to assess performance of the inference

Genome structures for simulations were randomly sampled 20Mb contiguous segments from the human genome. We included annotations for coding sequences (CDS), quiescent, and conserved noncoding regions acquired from GENCODE, ChromHMM, and PhastCons, respectively. The rest of the regions served as background noncoding regions. Sites within the top 5% of PhastCons470way scores were designated as conserved noncoding regions. We ran 20 replicates, each of which simulated the 20Mb segment sampled and output an SFS from 50 randomly drawn individuals (i.e. 100 chromosomes). Ten simulation replicates were randomly drawn and their respective SFSs were aggregated to generate a bootstrap SFS. We repeated this procedure ten times to obtain 10 bootstrap SFSs, each representing 200Mb of sequence.

We simulated a constant population size modeling the selection and neutral models using SLiM4.0.1 (Table 1, Haller & Messer 2019, 2023). In the selection model, all conserved noncoding sites and 20% of background noncoding sites were subject to negative selection. Selection coefficients for these mutations in these regions were drawn from a DFE with parameters derived from the inference performed on the top 5% of conserved sites in mammals (Table S2d). In the neutral model, all noncoding sites evolved neutrally. In both the selection and neutral models, nonsynonymous mutations are under selection and the synonymous mutations are neutral. The DFE parameters for nonsynonymous mutations were from the gamma distribution inferred from Kim et al., 2017 (the first row in the Table S4 in Kim et al., 2017 ). We divided the the population-scaled selection coefficient (2Nₐ*s*, shape=0.186, scale=875 ) by the ancestral population (N_anc_ = 12378, specified in the row5 (1KG, *μ*=1.5×10^-8^) in the Table S2 of Kim et al., 2017) to obtain the selection coefficient. The shape and scale parameters for the gamma distribution of selection coefficients for nonsynonymous mutations are 0.186, 0.0354. Table 1. Model and parameters for simulations.

**Table 1.** Model and parameters for simulations.

| Genomic regions | Selection model (DFE for heterozygote selection coefficient) | Neutral model |
| --- | --- | --- |
| exon: non-synonymous | shape=0.186,scale=0.0354 | Kim DFE |
| exon: synonymous | no selection | no selection |
| conserved noncoding | shape=0.1165,scale=0.005458 | no selection |
| quiescent | no selection | no selection |
| background | mix of 80% neutral and 20% conserved noncoding sites | no selection |

We then inferred demography and DFEs for the simulation replicates. For simulated data under a constant population size, we fit a two-epoch model with a single population size change. This model includes two parameters: *nu*, the ratio of the contemporary to the ancestral size, and *T*, the time of the size change (Fig.S11b). To simplify the sampling process from the simulations, we used all the quiescent sites for demography inference.

## DATA

### Functional annotation data

We download the ChromHMM states files from https://github.com/ernstlab/full_stack_ChromHMM_annotations and the Version 3 (May 6 2024 onwards) hg38 annotations from https://public.hoffman2.idre.ucla.edu/ernst/2K9RS/full_stack/full_stack_annotation_public_release/hg38 (Ernst & Kellis 2012; Vu & Ernst 2022). A more stable URL for ChromHMM data is https://zenodo.org/records/11015594. Each 200 bp window of the human genome is assigned to one of 100 chromatin states based on its epigenetic profile, which were previously grouped into biologically informative categories (Table_S1_ChromHMM_states: modified from Vu & Ernst 2022). The human Enhancers, Promoters, cis-enhancers, and distal-enhancers annotated by ENCODE are downloaded from SCREEN (https://screen.encodeproject.org/). We downloaded GENCODE annotation from http://ftp.ebi.ac.uk/pub/databases/gencode/Gencode_human/latest_release/gencode.v43.annotation.gtf.gz at 2023, July 10th (Frankish et al. 2022).

### Polymorphism data

We used 30x coverage NYGC 3202 samples from the 1000 Genomes Project sample collection ( https://www.internationalgenome.org/data-portal/data-collection/30x-grch38). Variants calling files from the 1000 genome project are downloaded from (http://ftp.1000genomes.ebi.ac.uk/vol1/ftp/data_collections/1000G_2504_high_coverage/working/202010 <u>28_3202_raw_GT_with_annot/</u>). We used polymorphisms from the Yoruba population from the 1KG project. We only include biallelic SNPs and high-quality sites that are included in the mask file (https://www.internationalgenome.org/announcements/genome-accessibility-masks/) on autosomes. The mask file was downloaded from http://ftp.1000genomes.ebi.ac.uk/vol1/ftp/data_collections/1000_genomes_project/working/20160622_ge <u>nome_mask_GRCh38/</u>.

### PhastCons scores

PhastCons17way and PhastCons470way scores were downloaded from https://hgdownload.soe.ucsc.edu/goldenPath/hg38/.

## Acknowledgements

We thank Kay Prüfer for helpful discussions about sequence data quality and Shamil Sunyaev for help with how to calculate the per site mutation rate from Roulette. CD, SR, and KEL were supported by the National Institutes of Health grant R35GM119856 to KEL.

## Extended data figures and tables Supplementary information

Supplementary Figures S1-S12.

Supplementary Tables: Table_S1_states_annotation.xlsx Table_S2_DFE.xlsx Table_S3_simulation.xlsx

