## Supplementary Figures for "The landscape of fitness effects of putatively functional noncoding mutations in humans"

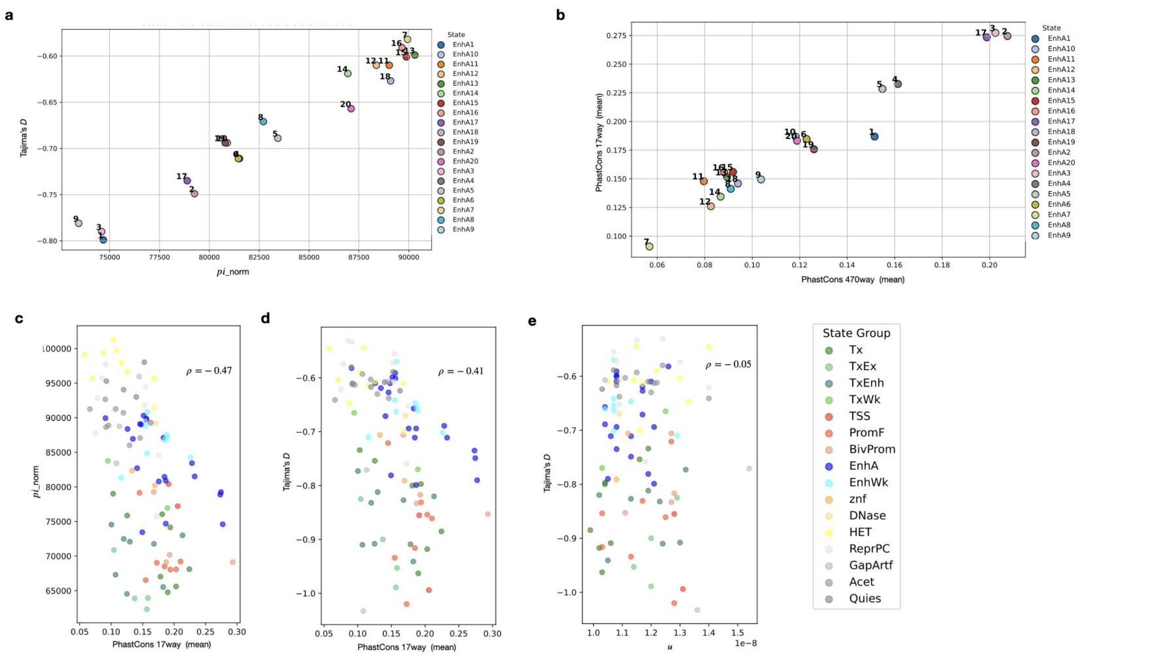


**Fig.S1: Patterns of genetic variation across chromatin state groups.**

**a,** Tajima's *D,* and *pi*_norm of each EnhA chromatin state. EnhA1: enhancers in most cell types, weaker in blood and enhancers in Digestive, Other, ES_deriv, Epithelial, Heart, Muscle, Thymus, and Neurosph; EnhA3: strong enhancers in most cells, weaker in blood and ESC&iPSC; EnhA9: Blood and thymus enhancer strongest; EnhA17: ESC, iPSC, Neurosph enhancers, and some differentiated; EnhA2: enhancers in mesenchymal, muscle, heart, neurosph, adipose. Additional information on EnhA states can be found in Table S1a. **b,** Mean of the PhastCons17way and PhastCons470way scores of each EnhA chromatin state. **c,** *pi*_norm correlates with PhastCons 17way scores with a Pearson's correlation coefficient of -0.47. **d,** Tajima's *D* correlates with PhastCons 17way scores with a Pearson's correlation coefficient of -0.41. **e,** Tajima's *D* is not correlated with the average haploid per-site per-generation mutation rates across chromatin state groups.

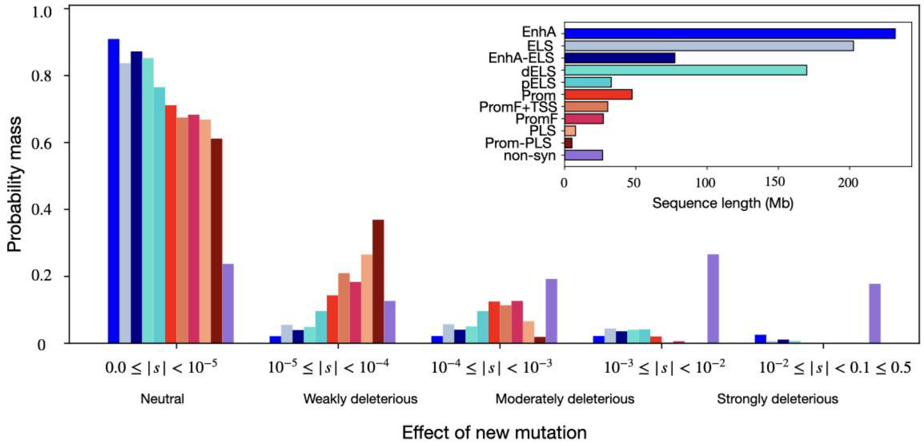


**Fig.S2: DFEs of mutations in different functional genomic regions.**

**a,** This figure presents results analogous to Fig.2a. EnhA denotes active enhancers annotated by ChromHMM, while ELS refers to enhancers annotated by ENCODE. EnhA-ELS represents the overlapping genomic regions between EnhA and ELS. dELS and pELS are distal and proximal enhancers, respectively, as defined by ENCODE. Prom includes ChromHMM states PromF, TSS, and BivProm. PromF+TSS combines the PromF and TSS ChromHMM states. PLS are promoters annotated by ENCODE, and Prom-PLS denotes genomic regions overlapping Prom and PLS. Coding regions are removed from all groups. Non-syn refers to non-synonymous mutations. b, The sequence length of conserved active enhancers (EnhA) and promoters (Prom) annotated by ChromHMM in Fig.2c and the number of non-synonymous sites.

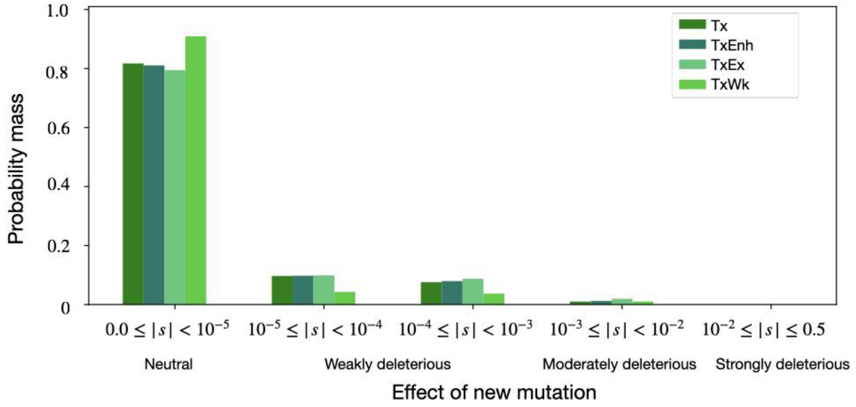


**Fig.S3: DFEs of transcribed genomic regions annotated by ChromHMM.**

The probability mass of new mutations across different ranges of selection coefficients ($|s|$) for non-coding transcribed genomic regions annotated by ChromHMM. Tx represents transcription; TxEnh, represents transcribed and enhancer; TxEx represents transcribed exon; and TxWk represents weak transcription.

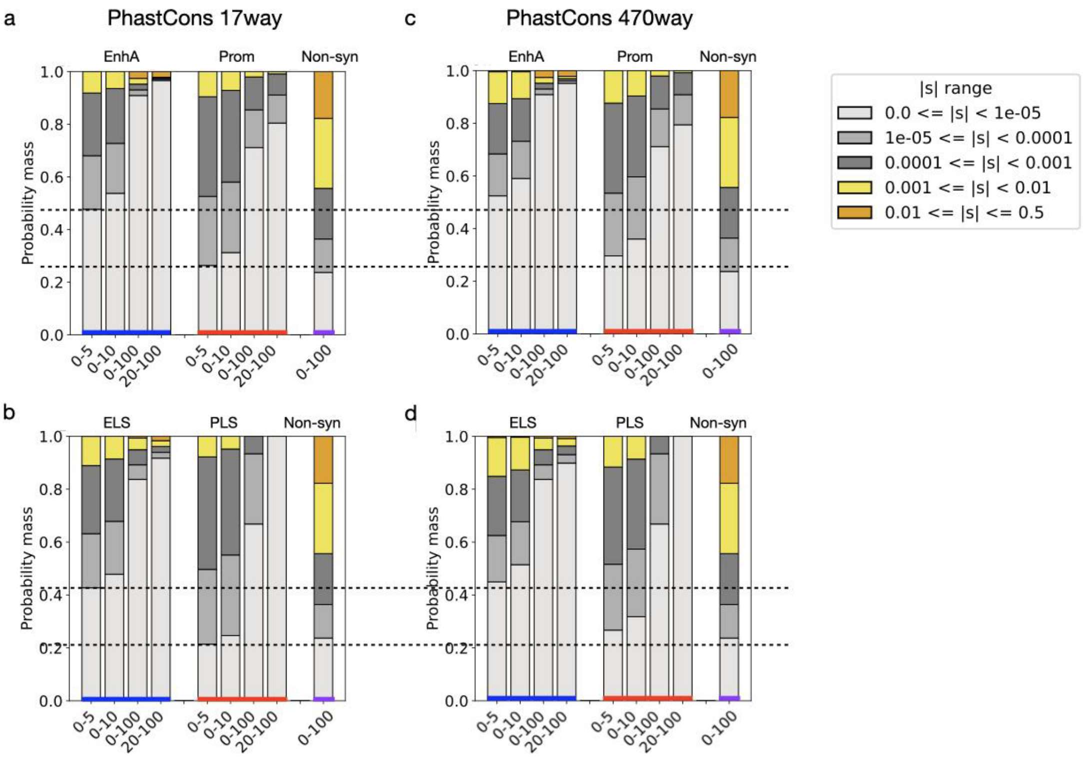


**Fig.S4: DFEs of conserved enhancers and promoters.**

Similar to Fig.2c, the DFEs of conserved enhancers and promoters. **a,b**, Conserved enhancers and promoters classified by topX% PhastCons 17way score. “EnhA” and “Prom” in (a) are enhancers and promoters annotated by ChromHMM. “ELS” and “PLS” in (b) are enhancers and promoters annotated by ENCODE. Non-syn indicates nonsynonymous mutations. **c,d,** Conserved enhancers and promoters classified by topX% PhastCons 470way score.

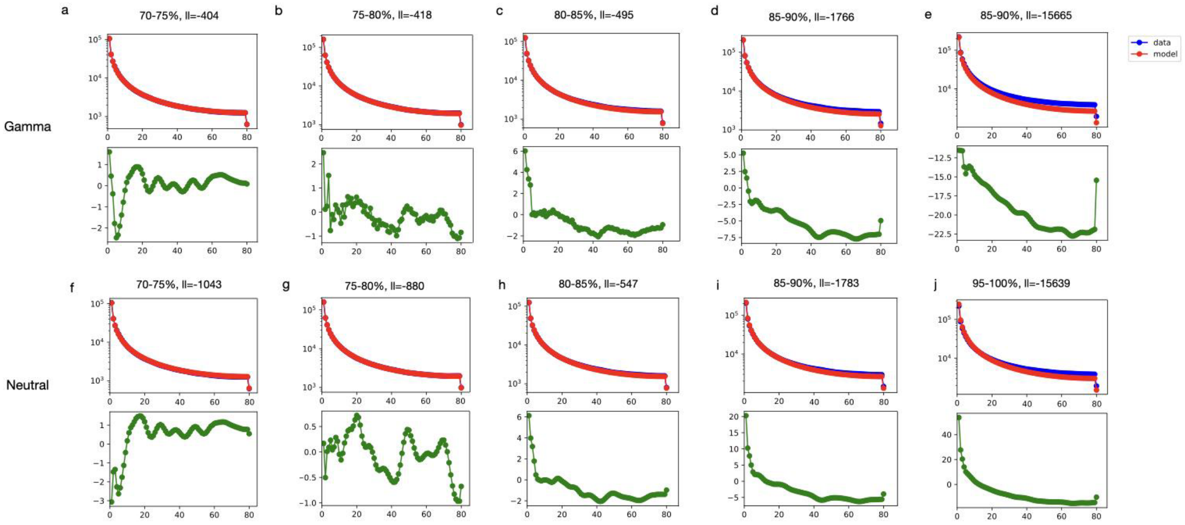


**Fig.S5. Difference in the SFS between the DFE models and data for candidate functional noncoding regions across different levels of constraint in primates.**

The site frequency spectra for the empirical data and best-fitting parameters as well as and residuals between the DFE models and data for candidate functional noncoding regions across different levels of constraint in primates with PhastCons17way scores smaller than the top 70%. **a-e**, The SFS and residuals of the gamma model. **f-j**, The SFS and residuals of the neutral model. For example, (a) represents the candidate functional noncoding regions with PhastCons17way scores ranging between 70-75%, and the log-likelihood (ll) of the gamma model is -404. The upper panel shows the number of alleles (y-axis) with given counts (x-axis), the bottom panel shows the normalized residual (residual = $(model - data)/\sqrt{model}$ ). Comparing the residual for the gamma model with the neutral model (f), the residuals of the gamma model are around zero, except for a few low-frequency alleles, while the residuals of the neutral model are larger. Moreover, the log-likelihood is much larger for the gamma model compared to the neutral model for candidate functional noncoding regions with PhastCons17way scores ranging between 70-75%. For candidate functional noncoding regions with PhastCons17way scores ranging between 75-80%, the residuals are both small for the gamma model (b) and the neutral model (g). For PhastCons17way scores ranging between 80-85%, residuals are almost the same between the gamma (c) and the neutral model (h). The log-likelihood difference is only a little larger than 50. For PhastCons17way scores ranging between 85-90%, the fit of the gamma (d) and the neutral (i) model is not as good as for the more constrained sites. For the last bin with the smallest PhastCons17way scores, although we prefer the neutral model (j), the fit of the neutral model is poor.

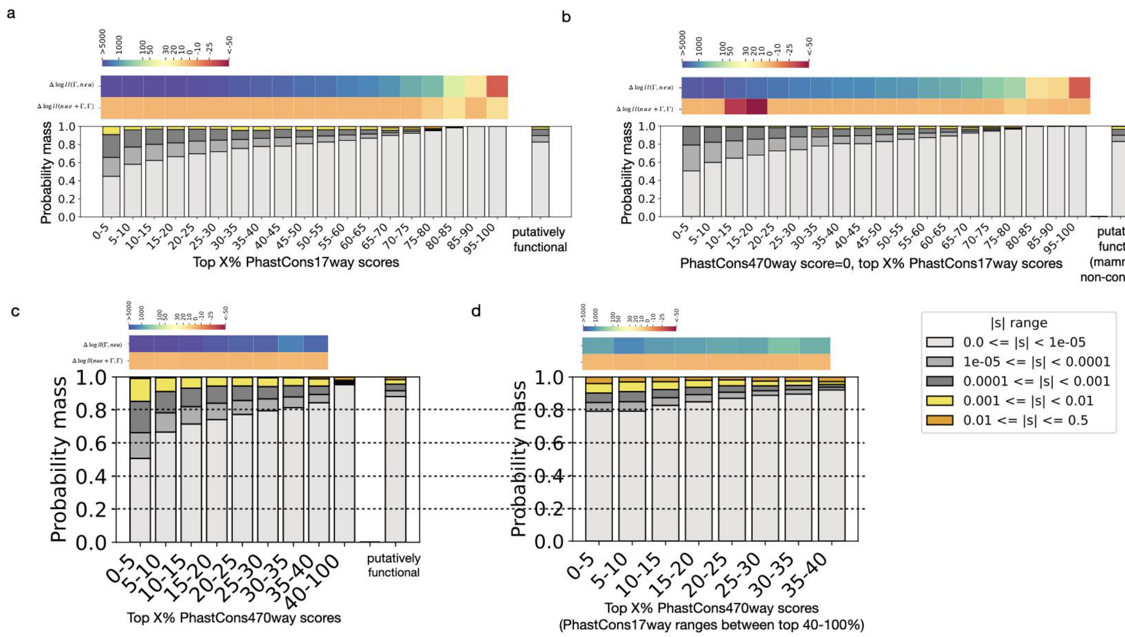


**Fig.S6: DFEs of candidate functional noncoding regions across different levels of constraint.**
a,b, Same as Fig.3a,b, but add the log-likelihood difference between the neugamma and gamma models. c, Similar to (a), but the conserved sites are categorized based on PhastCons 470way scores, and the bin of 40-100% includes a much larger group of sites. The DFE of the bin 40-100 is calculated as the weighted average of 12 randomly split groups for sites with a PhastCons 470way score of zero. Each group is weighted by the number of mutations (ul) it contains. d, Similar to (c) but limited to a subset of sites with PhastCons 17way scores in the bottom 60% and does not include the average DFE of putatively functional non-coding regions.

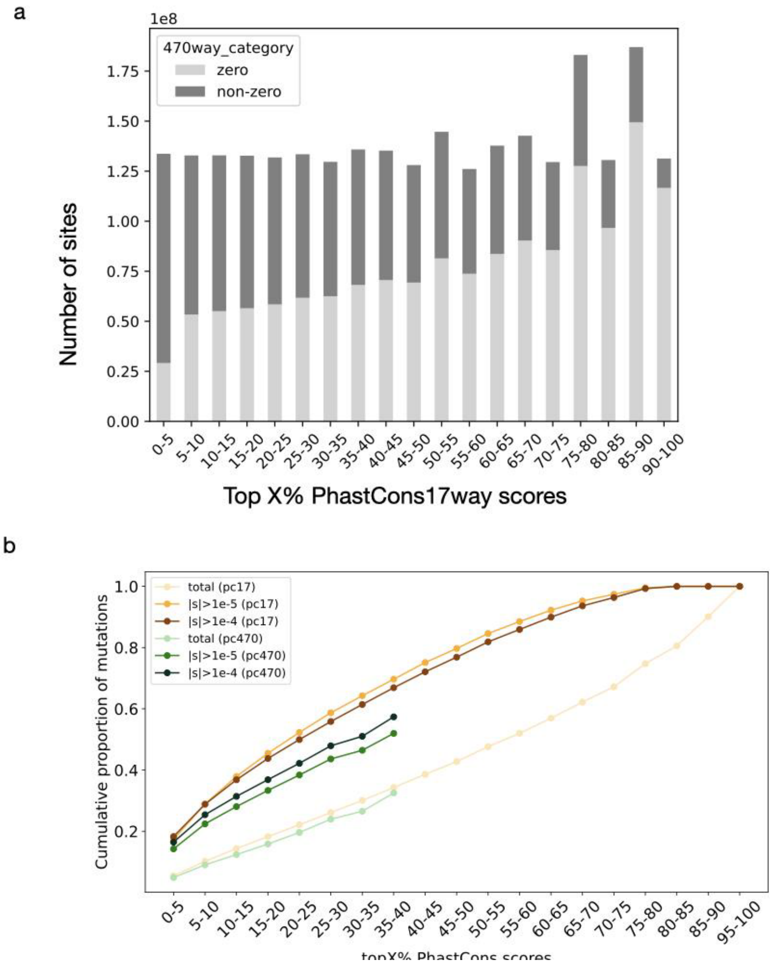


**Fig. S7. Number of sites and mutations across levels of conservation.**
**a,** Number of all sites—including putative functional noncoding genomic regions and other genomic regions with PhastCons scores—across bins of top X% PhastCons 17way scores, separated by mammalian conservation status. Light grey bars represent sites with a PhastCons 470way score of zero (i.e., not conserved across mammals), while dark grey bars represent sites with non-zero PhastCons 470way scores. **b,** Similar to Fig. 4a, cumulative proportions of total and non-neutral mutations across bins of PhastCons scores for putative functional noncoding genomic regions. The x-axis shows bins of top X% PhastCons scores. The orange line represents the cumulative proportion of non-neutral mutations ($|s|>10^{-5}$), the dark orange line represents the cumulative proportion of deleterious mutations ($|s|>10^{-4}$), and the light orange line represents all mutations in the top X% PhastCons 17way scores. The green line represents the cumulative proportion of non-neutral mutations ($|s|>10^{-5}$), the dark green line represents the cumulative proportion of deleterious mutations ($|s|>10^{-4}$), and the light green line shows all mutations in the top X% PhastCons 470way scores. The DFE of non-conserved regions in mammals is a weighted average of DFEs across bins of PhastCons 17way scores, with weights proportional to the number of mutations in each bin (see Methods: Binning conserved sites).

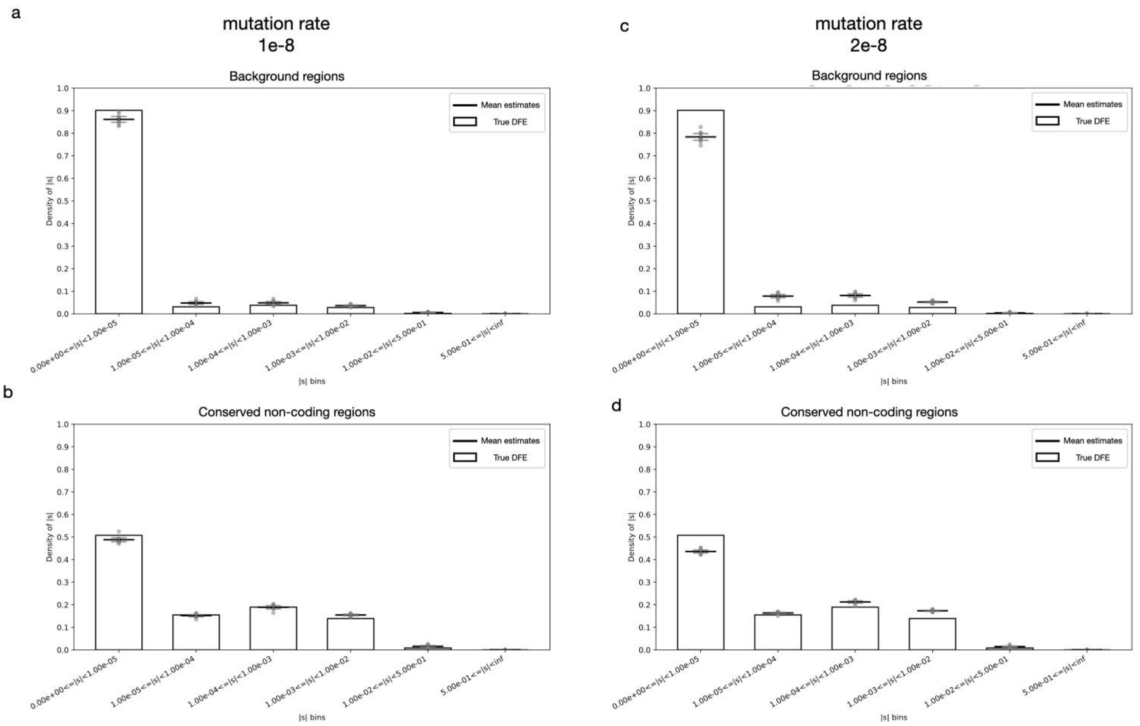


**Fig.S8: DFEs of simulations.**

**a,** DFE of background regions in simulations with a lower mutation rate. Bars indicate the true DFE. Grey points represent estimates from individual replicates, the long black horizontal line shows the mean of estimates, and short grey horizontal lines indicate the 95% confidence interval, assuming a Gaussian distribution (mean$\pm$standard deviation/$\sqrt{n}$, $n$=10).

**b,** Similar to (a), but showing the DFE of conserved non-coding regions.

**c,d,** DFEs of background **(c)** and conserved non-coding **(d)** regions from simulations with a higher mutation rate, following the same plotting scheme as in **(a)**.

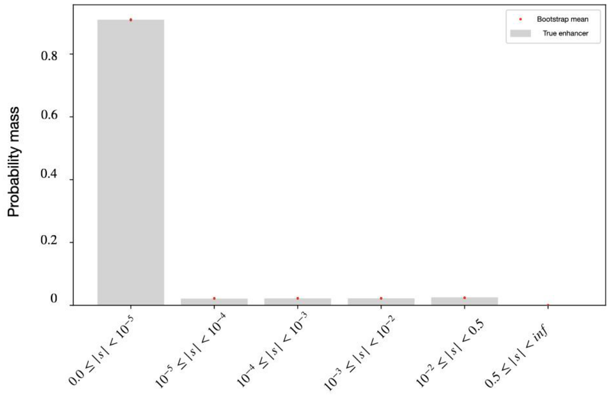


**Fig.S9: DFE of bootstrapped enhancers.**

DFEs of bootstrapped enhancers annotated by ChromHMM. The bars represent the DFE inferred from the real genome. The read point is the mean, and the tiny vertical line indicates the confidence interval. The confidence intervals are so small that they are almost invisible for deleterious mutations.

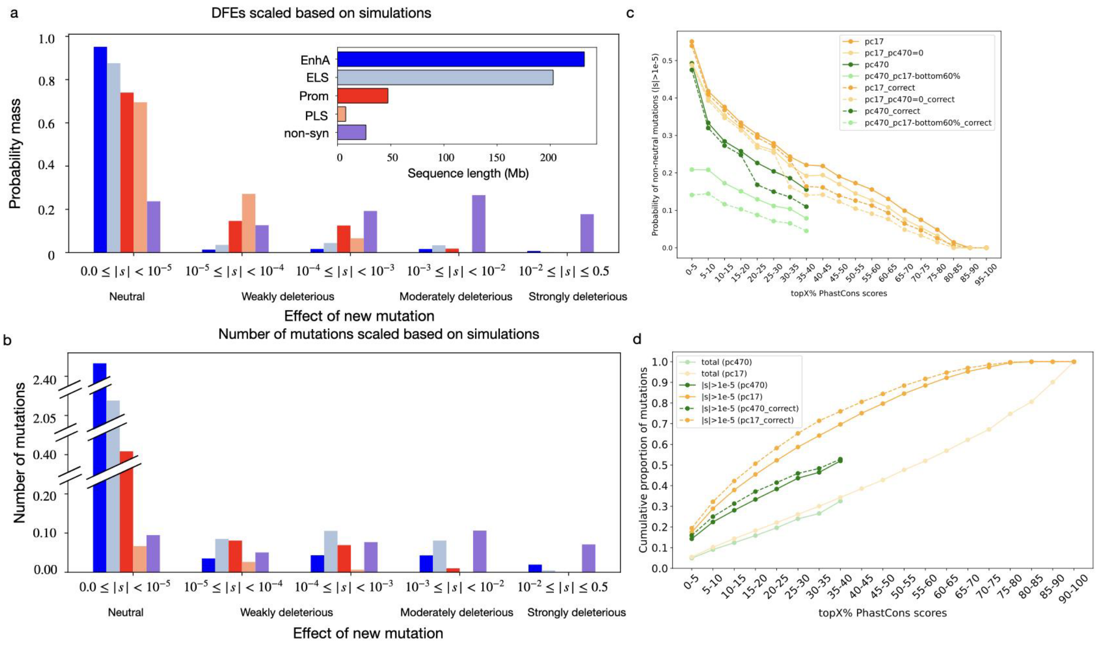


**Fig. S10. The DFE of mutations in enhancers and promoters after scaling based on simulations.**
**a,** Same as Fig.2a, but the DFEs are corrected based on the simulation results.
**b,** Similar to Fig.2b, but based on the corrected DFEs (dashed lines) in (a).
**c,** Same as Fig.3c but added dashed lines in the matched color showing the results based on corrected DFEs. For example, "pc17_correct" shows the proportion of non-neutral ($|s|>10^{-5}$) mutations across different levels of constraint based on PhastCons17way scores.
d, Cumulative proportions of total and non-neutral mutations across bins of PhastCons scores for putative functional genomic regions. The x-axis shows bins at the top X% PhastCons scores. The orange line is the same as in Fig.S6b, representing the cumulative proportion of non-neutral mutations ($|s|>10^{-5}$) in the top X% PhastCons 17way scores based on the originally inferred DFEs, the dashed orange line represents the cumulative proportion of non-neutral mutations ($|s|>10^{-5}$) based on the corrected DFEs according to simulations (Methods: Bias-corrected DFEs based on simulations), and the light orange line represents all mutations in the top X% PhastCons 17way scores. Similarly, the green line and the dashed green line represent the cumulative proportions of non-neutral mutations ($|s|>10^{-5}$) in the top X% PhastCons 470way scores based on the original and corrected DFEs, respectively. The light green line represents all mutations in the top X% PhastCons 470way scores.

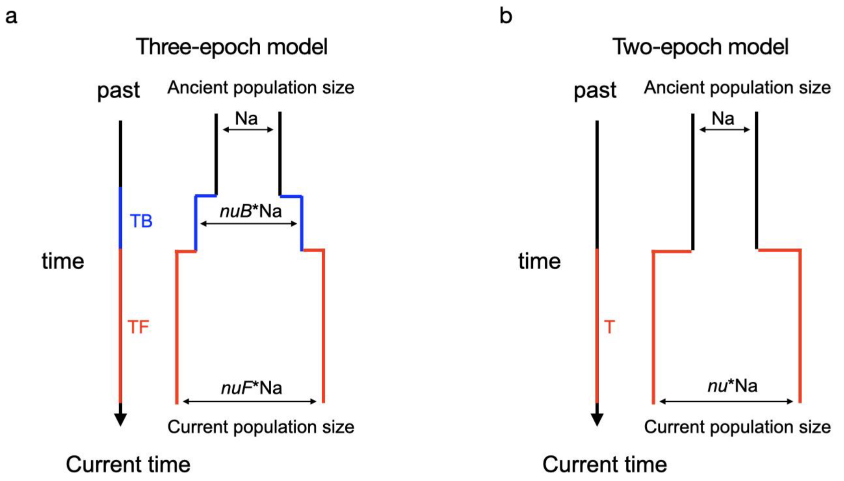


**Fig.S11: The demographic model.**
**a,** Three-epoch model. This model includes four parameters: the ratio of bottleneck to ancestral population size (nuB), the ratio of contemporary to ancestral size (nuF), the duration of the bottleneck (TB), and the time since recovery (TF). nuB and nuF are measured in the unit of ancestral population size, Na, and TB and TF are measured in units of 2×Na generations. nuB and nuF can take values greater than or less than one. **b,** Two-epoch model. This model includes two parameters: nu, the ratio of contemporary to ancestral size, and T, the time of the size change.

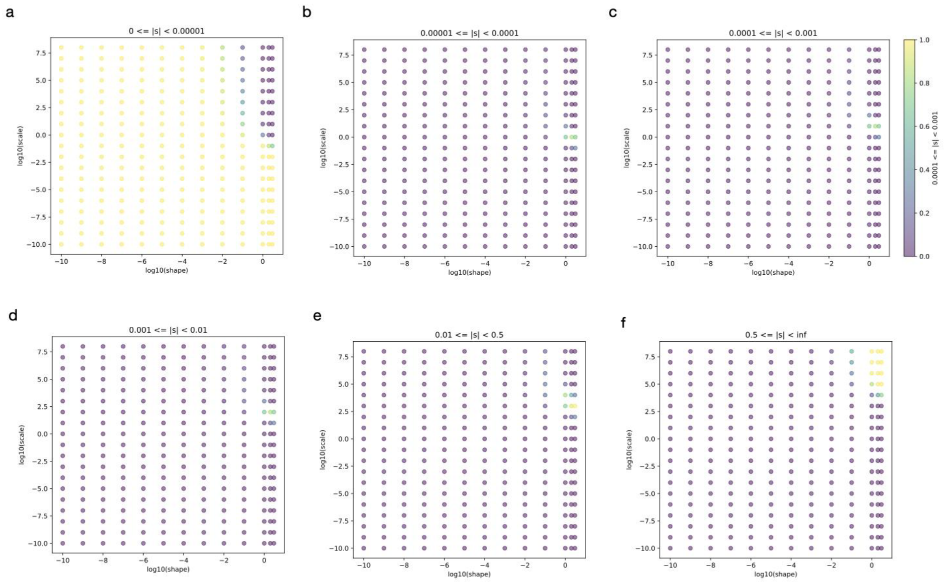


**Fig.S12: The landscape of the proportion of mutations of gamma models.**
The color indicates the proportion of mutations of a given range of selection coefficient (*s*, specified on the top left) of a gamma model of 2Na*s* with the given shape (x-axis) and scale parameters. The shape and scale parameters are log10-spaced.
